# Axon arrival times and physical occupancy establish visual projection neuron integration on developing dendrites in the *Drosophila* optic glomeruli

**DOI:** 10.1101/2022.08.20.504653

**Authors:** Brennan W. McFarland, HyoJong Jang, Natalie Smolin, Bryce W. Hina, Michael J. Parisi, Kristen C. Davis, Timothy J. Mosca, Tanja A. Godenschwege, Aljoscha Nern, Yerbol Z. Kurmangaliyev, Catherine R. von Reyn

**Affiliations:** School of Biomedical Engineering, Science and Health Systems, Drexel University, Philadelphia, PA; Department of Neuroscience, Vickie and Jack Farber Institute of Neuroscience, Thomas Jefferson University, Philadelphia, PA; Department of Biological Sciences, Florida Atlantic University, Boca Raton, FL; Janelia Research Campus, Howard Hughes Medical Institute, Ashburn, VA; Department of Biology, Brandeis University, Waltham, MA; Department of Neurobiology and Anatomy, Drexel University College of Medicine, Philadelphia, PA

**Keywords:** Pupal Development, *Drosophila melanogaster*, Giant Fiber, Optic Glomeruli, Neural Activity, Adaptation, Electrophysiology, Visual Projection Neurons, Visual System, Descending Neuron

## Abstract

Behaviorally relevant, higher order representations of an animal’s environment are built from the convergence of visual features encoded in the early stages of visual processing. Although developmental mechanisms that generate feature encoding channels in early visual circuits have been uncovered, relatively little is known about the mechanisms that direct feature convergence to enable appropriate integration into downstream circuits. Here we explore the development of a collision detection sensorimotor circuit in *Drosophila melanogaster*, the convergence of visual projection neurons (VPNs) onto the dendrites of a large descending neuron, the giant fiber (GF). We find VPNs encoding different visual features establish their respective territories on GF dendrites through sequential axon arrival during development. Physical occupancy, but not developmental activity, is important to maintain territories. Ablation of one VPN results in the expansion of remaining VPN territories and functional compensation that enables the GF to retain responses to ethologically relevant visual stimuli. GF developmental activity, observed using a pupal electrophysiology preparation, appears after VPN territories are established, and likely contributes to later stages of synapse assembly and refinement. Our data highlight temporal mechanisms for visual feature convergence and promote the GF circuit and the *Drosophila* optic glomeruli, where VPN to GF connectivity resides, as a powerful developmental model for investigating complex wiring programs and developmental plasticity.

## INTRODUCTION

In a developing brain, the coordinated wiring of multiple inputs onto a neuron and organization of these inputs across a neuron’s dendrites establish the computational role for that neuron. Uncovering the mechanisms that assemble and localize multiple inputs is pivotal to understand how inputs are miswired in neurodevelopmental disorders^1-3^ and how developmental processes attempt to compensate when particular inputs are missing or fail to connect^4,5^. Across species, we know little about how multiple inputs that converge upon a neuron are wired during development because the underlying circuits are often not well established – we are missing the solution to the wiring program, where all inputs are known and synapse locations are mapped.

Here, we capitalize on recent connectome data and functional investigations within the *Drosophila* optic glomeruli, a central brain region where visual feature inputs converge onto sensorimotor circuits^6-10^. Optic glomeruli are the output region for columnar visual projection neurons (VPNs) that are hypothesized to encode visual features^9,11-14^. VPN dendrites are retinotopically distributed to tile the lobula and the lobula plate of the fly’s optic lobes, while fasciculated VPN axons terminate within their respective glomerulus^11^. Within each glomerulus, VPNs synapse with multiple targets, including descending neurons (DNs) that project axons to the ventral nerve cord (VNC, the fly spinal cord homologue) where they in turn synapse onto interneurons and motoneurons that generate behavioral outputs^15-17^. Unlike the *Drosophila* olfactory glomeruli which have a predominantly one to one olfactory receptor neuron to projection neuron mapping^18^, each VPN glomerulus is not dedicated to a single DN type. Instead, DN dendrites infiltrate multiple, semi-overlapping subsets of glomeruli^10,16,19^, essentially assembling VPN features into higher order, behaviorally relevant motor and premotor outputs. How this complex wiring of visual feature inputs onto DNs is established during development is presently unknown.

A pair of large DNs called the giant fibers (GFs) receive major input from two VPN types in the optic glomeruli: lobula columnar type 4 (LC4) neurons and lobula plate - lobula columnar type 2 (LPLC2) neurons (Figure 1A)^6,7,13^. LC4 and LPLC2 encode the angular velocity and angular size of an expanding object, respectively^6,7^, and enable the GFs to drive a rapid takeoff escape in response to an object approaching on a direct collision course^17^. The GF circuit within the optic glomeruli presents an ideal model for developmental investigations. GF connectivity in adult flies has been recently established through electron microscopy, genetic access exists for the GF and its major visual input cell types, and the large GF dendrites within the glomeruli can be resolved and tracked across development^6,7,20,21^. Additionally, the accessibility of the GF to electrophysiology enables the functional consequences of developmental events to be directly evaluated^6,7^. GF dendrites are also in close proximity to VPNs that are not synaptic partners in the adult, like lobula plate - lobula columnar type 1 (LPLC1) neurons. This provides an opportunity to investigate developmental interactions with cell types that do or do not select the GF as a synaptic partner.

**Figure 1.**
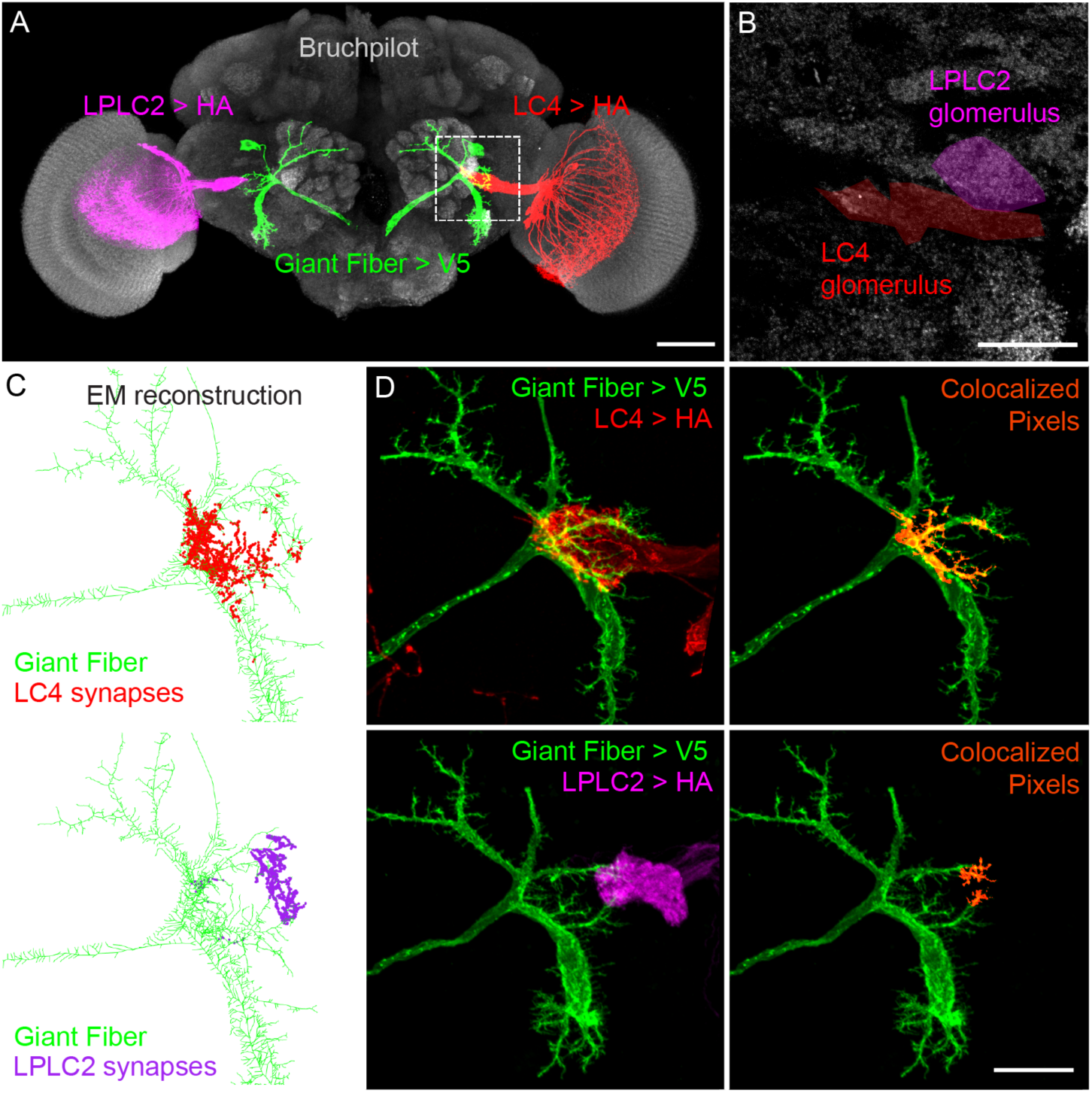
LC4 and LPLC2 occupy distinct regions on GF dendrites. **(A)** GF (green), LPLC2 (magenta, one hemisphere), and LC4 (red, one hemisphere) maximum intensity projections superimposed over neuropil label Bruchpilot (Brp, gray) Scale bar, 50μm. **(B)** Optic glomeruli as identified by Brp labeling with the LPLC2 (magenta) and LC4 (red) glomeruli highlighted. Maximum intensity projection of a substack located within the dashed box in (***A***). Scale bar, 20μm. **(C)** *Drosophila* hemibrain EM reconstruction of GF (green) with colored dots indicating synapses from LC4 (red, top) and LPLC2 (magenta, bottom). **(D)** (Left) Maximum intensity projections of dual labeled GF and VPNs. (Right) Colocalized pixels (orange) between GF and respective VPNs superimposed over GF maximum intensity projections. Scale bar, 20μm.

Here, we establish the GF circuit^6,7,17^ as a model for visual feature convergence in a developing nervous system. We screened VPN and GF GAL4 and LexA driver lines for early developmental expression and cell-type specificity. We then used identified driver lines to characterize the timecourse of VPN and GF interactions across metamorphosis that lead to their final organization in the adult. Combining a comprehensive single-cell RNA sequencing (scRNA-seq) atlas of the developing *Drosophila* visual system^22^, synaptic protein labeling over development, and a novel *ex-plant* electrophysiology preparation that enabled us to record from the GF at distinct developmental timepoints, we correlated the time course of VPN to GF interactions with the arrival of synaptic machinery and neural activity. To determine how competition shapes VPN synapse organization along GF dendrites, we genetically ablated one VPN cell type (LC4) and investigated both structural and functional compensation from the surviving VPN partner (LPLC2). Our data provide a thorough characterization of the assembly of visual feature convergence onto GF dendrites and establish the optic glomeruli as a genetically and functionally tractable model to uncover mechanisms underlying complex wiring programs.

## RESULTS

### VPNs are localized to stereotyped regions on GF dendrites

GF dendrites extend into the optic glomeruli in close proximity to multiple VPN cell-types (Figure 1A,B). EM reconstruction of a full adult fly brain (FAFB^20^) previously revealed that 55 LC4 and 108 LPLC2 neurons connect directly onto GF optic glomeruli dendrites, contributing 2,442 and 1,366 synapses, respectively^7^. VPN synapses segregate across the medial-lateral axis, with LC4 predominantly localized to medial, and LPLC2 to lateral, dendritic regions^7^. To investigate stereotypy in VPN to GF connectivity, we utilized a second EM dataset of a *Drosophila* hemibrain^21,23^. In this EM reconstruction, we found LC4 (71/71) and LPLC2 (85/85) neurons established 2,290 and 1,443 synapses onto the GF optic glomeruli dendrites, respectively (Figure 1C). We confirmed LC4 and LPLC2 synapses segregate along the medial-lateral axis, with only 2/85 LPLC2 neurons making synapses in predominantly LC4 occupied medial areas.

Since existing EM data only represent connectivity within two fly brains, we further investigated localization stereotypy by examining contacts between GF and VPN membranes across multiple adult flies. We used split-GAL4 driver lines that selectively labeled LC4 (*LC4_4- split-GAL4*, generated for this paper) or LPLC2 (*LPLC2-split-GAL4*)^11^, and simultaneously labeled the GF with a LexA line (*GF_1-LexA)^24^* (Figure 1D, left). As a proxy for membrane contacts, we performed intensity-based thresholding on each cell-type of interest to generate representative masks, and then visualized colocalized regions along GF dendrites (Figure 1D, right). We consistently observed LC4 contacts on the most medial regions of the GF optic glomeruli dendrites and LPLC2 contacts on the most lateral regions. We also found on occasion (4/17 brain hemispheres), as seen in the hemibrain dataset, a small subset of LPLC2 axons extending into the most medial regions on the GF (Supplemental Figure 1, arrow)^21^. These data suggest that LC4 and LPLC2 consistently segregate to stereotyped regions along the medial-lateral axis with rare exceptions.

We used the same approach to assess the projections of LPLC1 (*LPLC1_1-split-GAL4)^11^*, a cell-type adjacent to LC4 and LPLC2 that does not synapse directly with the GF^7^. As expected, no synapses were identified in the hemibrain EM dataset (Supplemental Figure 2) and no membrane contacts were observed between GF and LPLC1 across all adult flies imaged with confocal microscopy (Supplemental Figure 2). We additionally employed GFP reconstitution across synaptic partners (GRASP)^25,26^ to visualize contacts between adjacent membranes and observed GFP expression between GF and LC4/LPLC2 in their respective medial/lateral locations, but not between GF and LPLC1 (Supplemental Figure 3).

### GF lateral dendrites extend, elaborate, and then refine across pupal stages

Following our detailed anatomical characterization, we sought to determine how the precise VPN localization along GF dendrites arises across development. Prior developmental investigations into the GF have focused on axonal wiring with respect to postsynaptic interneuron and motor neuron partners in the ventral nerve cord (VNC)^27-32^. However, little is known about how GF dendrites develop in the central brain^33^. The GF is born during embryonic stages but does not generate neurites until the third instar larval stage^27^. To track the GF at these early timepoints, we used the *GF_1-LexA* line that labels the GF starting in late larval stages (Supplemental Figure 4) and dissected pupae in 12-hour increments over metamorphosis, a period marked as the time between pupa formation and eclosion.

Across development, we tracked the complexity and size of GF optic glomeruli dendrites by quantifying their volume (Figure 2A,B) and the length of the maximum dendrite extension along the medial-lateral axis (Figure 2A,C). In the early stages of metamorphosis, 24-48 hours after pupa formation (hAPF), the GF exhibited numerous filopodia, long thin protrusions without a bulbous head, and arbor complexity increased with the GF projecting between 3.1 ± 1.0 primary dendrites laterally (Figure 2A,B). During the middle stages of metamorphosis, from 48 hAPF to 60 hAPF, the GF dendrites had the largest increase in their medial-lateral extent (Figure 2C), followed by a peak in the overall volume and extension length at 72 hAPF (Figure 2A-C). During this time, filopodia were still present, but visibly shorter than in the first half of metamorphosis. In the final stages of metamorphosis from 72 hAPF to eclosion, the volume of GF dendrites significantly decreased (Figure 2A-C), while the medial-lateral length was maintained. Filopodia were no longer obvious, and branches appeared less complex and began to resemble their adult morphology.

**Figure 2.**
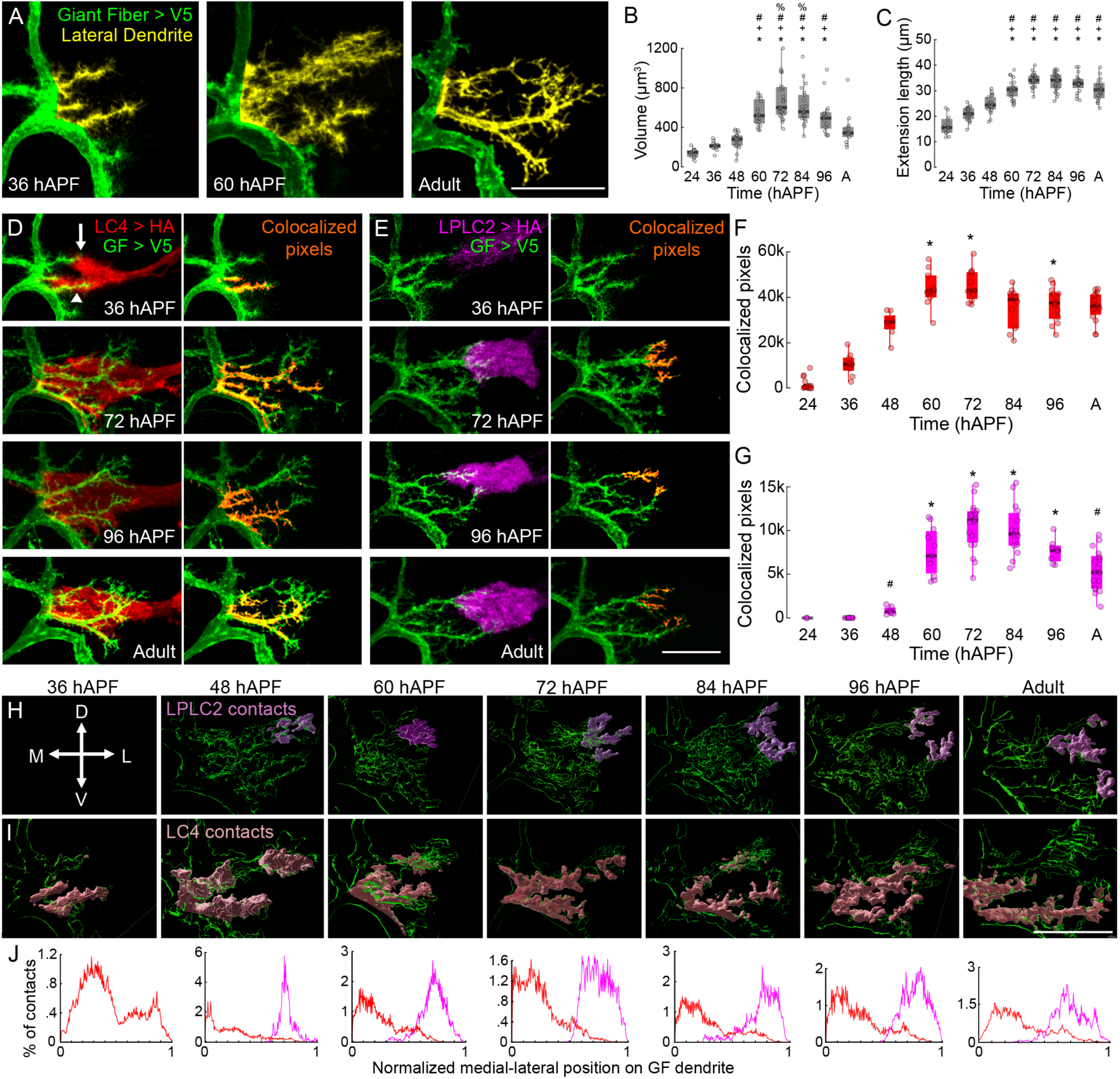
LC4 and LPLC2 territories on GF dendrites are established early in development. (**A**) Maximum intensity projections of GF (green) 36 hAPF (left), 60 hAPF (middle), and in adult (right) with the VPN dendritic region highlighted in yellow at distinct developmental stages. Scale bar, 20μm. (**B**) Quantification GF lateral dendrite volume from (A). Unpaired Kruskal-Wallis test (p = 1.339 x 10^−18^), Tukey-Kramer multiple comparison test post hoc, * = p < .05 as compared to 24 hAPF, + = p < .05 as compared to 36 hAPF, # = p < .05 as compared to 48 hAPF, and % = p < .05 as compared to adult. N ≥ 13 hemibrains from ≥ 10 flies. (**C**) Quantification of maximum dendrite extension length across the medial-lateral axis. Unpaired Kruskal-Wallis test (p = 2.072 x 10^−12^), Tukey-Kramer multiple comparison test post hoc, * = p < .05 compared to 24 hAPF, + = p < .05 compared to 36 hAPF, # = p < .05 compared to 48 hAPF. **(D,E)** Left, maximum intensity projections of GF (green) with respect to LC4 (red, D), and LPLC2 (magenta, E) axonal membrane at distinct developmental stages. Right, maximum intensity projections of GF with VPN colocalized pixels (orange) superimposed along GF dendrites. Arow and arrowheads indicate divergent dorsal and ventral VPN axons, respectively. Scale bar, 20μm. **(F,G)** Quantification of colocalization in (D,E) with colors corresponding to VPN type. Unpaired Kruskal-Wallis test (LC4, p = 2.088 x 10^−12^; LPLC2, p = 1.983 x 10^−10^), Tukey-Kramer multiple comparison test post hoc, * = p < .05 as compared to 36 hAPF, + = p < .05 as compared to 60 hAPF, # = p < .05 as compared to 72 hAPF. N ≥ 6 hemibrains from ≥ 3 flies. **(H,I)** 3D renderings of GF lateral dendrites (green) with LPLC2 (H, magenta) or LC4 (I, red) colocalized pixels superimposed at distinct timepoints during development. Scale bar, 20μm. D - dorsal, V – ventral, M – medial, L – lateral. **(J)** Histograms of the spatial distribution of LC4 and LPLC2 contacts along the normalized medial-lateral GF dendrite axis across development; colors are the same as in (H, I). N are as stated in (F,G).

### Initial contacts between GF and VPNs are staggered in time

We next investigated VPN axon targeting with respect to GF dendritic outgrowth. At present, it is unknown when columnar VPN neurons are born^34^, but these neurons may arise in late larval to early pupal stages, a period when neuroblasts give rise to visual neurons (such as T4/T5) that provide input to VPN dendrites in the lobula and lobula plate^35-37^. We hypothesized VPNs would commence outgrowth and partner matching in coordination with GF dendrite development. To visualize developmental interactions of select VPN and GF, we used existing^11^ or newly developed VPN split-GAL4 driver lines screened for pupal expression (Supplemental Figure 4), to concurrently label select VPN cell-types (*LC4_4-split-GAL4*, *LPLC2-split-GAL4, LPLC1_1-split-GAL4*) and the GF (*GF_1-LexA*) over metamorphosis. To quantify interactions, we employed our membrane colocalization method (Figure 1D) instead of synapse labeling methods (such as t-GRASP^38^) because we wanted to track all putative interactions, including those that precede synapse formation, over time and did not want to create ectopic adhesions between membranes. We additionally compared our membrane colocalization method to a GRASP variant that is not restricted to presynaptic terminals^25^ and found no statistical difference between the number of colocalized or GFP positive pixels (Supplemental Figure 5).

Using colocalization as a proxy for membrane contacts, we investigated developmental interactions between GF and its known adult partner VPNs, LC4 and LPLC2. We observed LC4 axonal extension and initial contact with GF at 24 hAPF (Figure 2F). At this timepoint, LC4 axons diverge into a dorsal fraction projected near the dorsal branch of the GF optic glomeruli dendrites, and a ventral fraction projected towards the proximal regions of the GF dendrites (Supplemental Figure 6). At 36 hAPF, contacts between the GF and LC4 increased, with both dorsal and ventral fractions still apparent (Figure 2D, arrow, arrowhead, respectively). Although *LPLC2-split-GAL4* shows obvious expression at this time, we did not observe any contacts between LPLC2 and the GF (Figure 2E,G).

At 48 hAPF, LC4 and GF continued to show an increase in contacts (Figure 2F), and LC4 dorsal and ventral axons had converged (Supplemental Figure 6). At this time, approximately 24 hours after initial GF and LC4 contact, we observed GF contacts with LPLC2 (Figure 2E,G) as the dorsal branch of the GF dendrites extended past LC4 axons (Supplemental Figure 6, arrowhead). Altogether, our data suggest that during the first half of metamorphosis, as the GF is seeking out synaptic partners, interactions with VPN are staggered in time.

After the initial establishment of contacts, we next observed a significant increase in contacts between partner VPNs and GF from 60 hAPF to 72 hAPF (Figure 2D-G). At 84 hAPF through eclosion, contacts between GF and both VPNs decreased and then stabilized. Our results suggest that in the second half of metamorphosis, GF prioritizes dendritic outgrowth, enhances contacts with partner VPN, and eventually refines and stabilizes contacts with appropriate VPN partners, LC4 and LPLC2.

We next investigated interactions between GF and a neighboring VPN, LPLC1, that does not maintain synapses with GF in adulthood (Supplemental Figure 2). We observed a relatively small number of contacts between GF and neurons labeled with *LPLC1_1-split-GAL4*^11^ appearing around 48 hAPF, peaking around 60 hAPF, and disappearing around 84 hAPF (Supplemental Figure 7A,B). This driver line, however, may also label a subset of VPN that are not LPLC1 during development (Supplemental Figure 7D,E), so we repeated our contact analysis by generating two new LPLC1 driver lines, *LPLC1_2-split-GAL4* and *LPLC1_3-split-GAL4* (Supplemental Figure 4). While these driver lines revealed GF and LPLC1 membranes are adjacent at 60 hAPF, we observed minimal to no contacts with GF (Supplemental Figure 7C).

Altogether, these results suggest that early in development, GF contacts are already biased towards VPNs that are synaptically coupled to the GF in the adult.

### LC4 and LPLC2 occupy and maintain distinct regions along the GF dendrite

In the adult GF circuit, LC4 inputs are localized to the medial regions of GF dendrites, whereas LPLC2 inputs are localized to the most lateral regions (Figure 1). It is unknown if this medial-lateral segregation is established initially or arises over development. To address this, we manually aligned the GF dendrites across brains and quantified the density of contacts along the medial-lateral axis. Across all time points, we found minimal overlap between LC4 and LPLC2; the peak density of contacts for LC4 and LPLC2 consistently occupied the most medial and lateral regions, respectively (Figure 2H-J).

The GF optic glomeruli dendrites contain dorsal and ventral branches, therefore we repeated the analysis of membrane contacts along the dorsal-ventral axis. At 48 hAPF, LPLC2 contacts were primarily confined to the dorsal regions, and LC4 to the ventral regions (Supplemental Figure 8). This segregation was reduced at 60 hAPF and less obvious in the later stages of development (84 hAPF – 96 hAPF), in alignment with previous investigations into adult synapse localization along the dorsal-ventral axis^7^. Altogether, our results highlight the importance of the medial-lateral division of LC4 and LPLC2 inputs onto GF dendrites, where targeting is established early and maintained throughout development.

### Upregulation of synaptic machinery across key stages of metamorphosis

Although our colocalization data provide the time course for interactions between VPN axons and GF dendrites across development, they do not provide information on the timing of synaptogenesis. We therefore investigated how our time course for GF/VPN interactions aligned with the expression of presynaptic machinery (Figure 3A,B). We used a comprehensive scRNA-seq atlas of the developing *Drosophila* visual system which profiled optic lobe neurons at multiple time points across metamorphosis and identified both global and cell-type specific transcriptional programs^22^.

**Figure 3.**
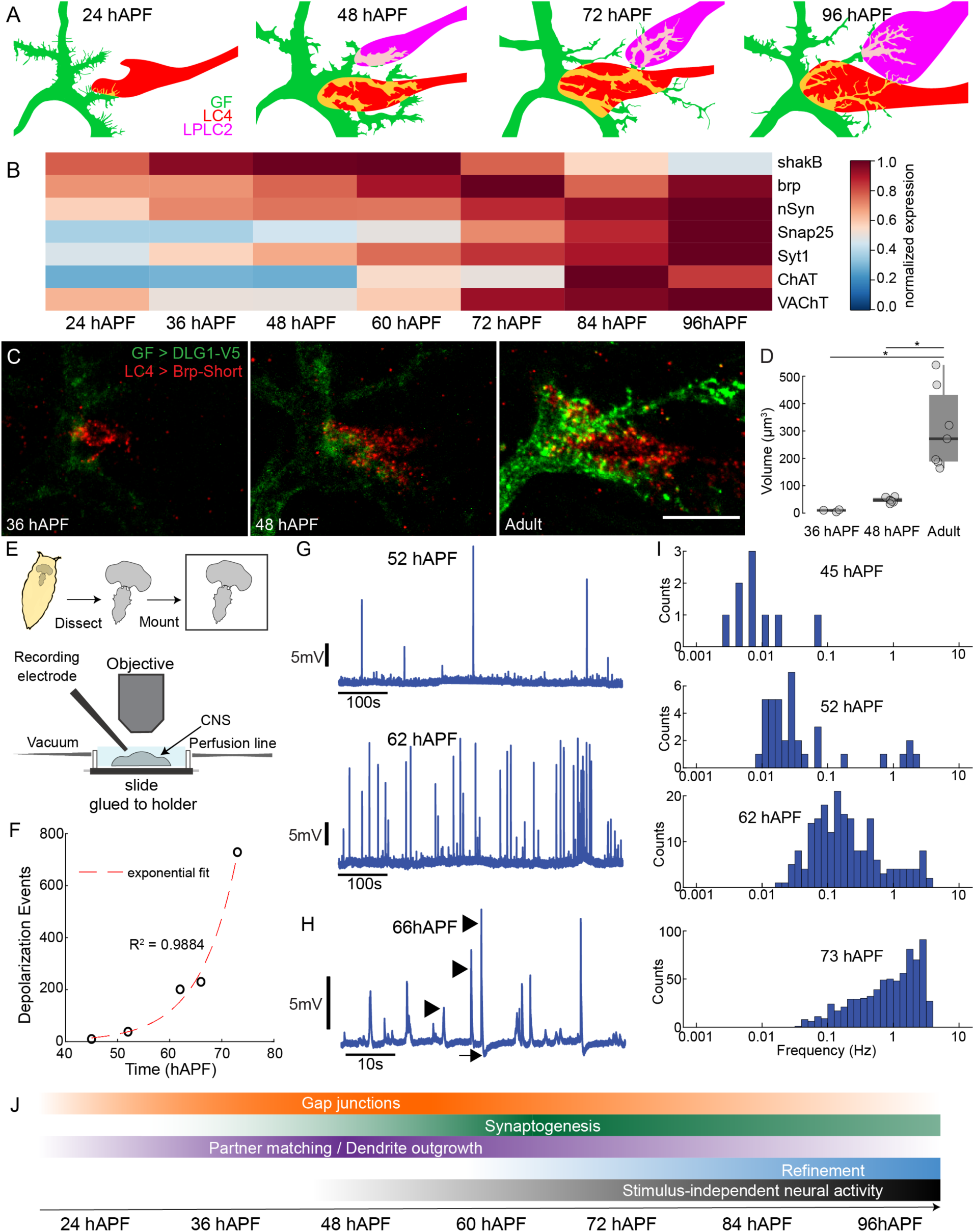
Synaptogenesis and the emergence of stimulus-independent neural activity. **(A)** Schematic of GF and VPN developmental interactions **(B)** Heatmap timecourse of average, normalized VPN mRNA expression of genes for electrical and chemical synapse function. Data are from the optic lobe transcriptional atlas^22^ and individual VPN expression patterns can be found in Supplemental Figure 9. **(C)**Max intensity projections of a substack of DLG expression in GF (DLG1-V5, green) and Brp puncta in LC4 (Brp-Short, red) at selected timepoints. Scale bar, 10μm. **(D)**Quantification of volume of Brp colocalized with DLG from (C). Unpaired Kruskal-Wallis test (p = 0.001), Dunn-Sidak comparison test post hoc, * = p < .05, N= 3-7 hemibrains from 2-5 flies. **(E)**Schematic of *ex-plant* pupal electrophysiology preparation. **(F)**The total number of identified depolarizing events increases exponentially (fit, dotted red line) over time. N = 5 flies. **(G)**Representative traces of GF membrane potential recordings using the pupal electrophysiology preparation for two timepoints. **(H)**Zoomed in recording showing features resolvable with electrophysiology. Arrow indicates hyperpolarization following large depolarizing events, arrowheads indicate different event amplitudes. **(I)**Distribution of event frequencies from inter-event intervals. **(J)**Timecourse of developmental stages as estimated from anatomical, scRNA-seq and electrophysiology data.

As developmental clusters corresponding to LC4, LPLC2, and LPLC1 have been identified in this dataset, we re-analyzed these data to determine when genes required for synaptic transmission were upregulated in metamorphosis. We first investigated the expression of *brp*, a presynaptic active zone protein that is homologous to the mammalian ELKS/CAST family^39,40^ that is commonly used to label presynaptic terminals. We found *brp* to be present as early as 24 hAPF and gradually increase up until 60 – 72 hAPF (Figure 3B, Supplemental Figures 9,10). Presynaptic genes in the SNARE complex^41-44^ *nSyb*, *cpx*, *Snap25* and *Syx1A* were also present at early pupal stages, but significant upregulation was delayed with respect to *brp*, from around 60 hAPF until the end of metamorphosis (Figure 3B, Supplemental Figures 9,10).

LC4, LPLC1 and LPLC2 are predicted to be cholinergic^44^, and we found genes for cholinergic synapse function (*ChAT* and *VAChT*) were upregulated in the late stages of metamorphosis (≥ 60 hAPF) (Figure 3A, Supplemental Figure 9-10), delayed from the initial appearance of presynaptic machinery, but following the time course reported for other cholinergic neurons^22^. This upregulation coincides with our observed decrease in GF dendritic complexity, and refinement and stabilization of GF and VPN contacts (Figure 2F,G, Supplemental Figure 7A-C). These data suggest that although a subset of presynaptic components are expressed and potentially assembled early, VPN cholinergic machinery arrives too late to contribute to the initial targeting and localization of VPN axons on GF dendrites. Cholinergic activity instead is likely to participate in VPN and GF synapse refinement and stabilization.

Given the significant role electrical synapses play across development, we also examined expression of the innexin family of gap junction proteins. Shaking-B (*shakB*) has been denoted to be the predominant innexin over development^22^, and we also observed a significant increase as early as 36 hAPF (Figure 3B, Supplemental Figures 9,10) with all other innexins showing minimal to no expression across metamorphosis. *shakB* expression peaked between 48-60 hAPF, followed by a significant decrease from 60 hAPF to 72 hAPF for all cell types (Supplemental Figures 9,10). Interestingly, this decrease occurred as *ChAT* and *VAChT* increased, potentially marking the transition from predominantly electrical synaptic coupling to chemical synaptic coupling. A summary of our scRNA-seq analyses aligned to our developmental interaction timecourse can be seen in Figure 3A,B.

### GF and VPN synapse assembly is initiated during the partner matching stages

While our scRNA-seq data provide an estimate of gene expression across development, relative levels of mRNA do not necessarily correlate linearly to protein translation^45^. Therefore, using our scRNA-seq data to guide our hypotheses, we next investigated the temporal expression patterns of select pre- and postsynaptic proteins in VPN and GF. We utilized an iteration of Synaptic Tagging with Recombination (STaR)^46^ to visualize LPLC2 and LC4 specific Brp, driven by its endogenous promoter and tagged with smGdP-V5. From our scRNA-seq data, *brp* is expressed early in both LC4 and LPLC2 (Supplemental Figures 9,10). Using *LC4_4-split-GAL4* and *LPLC2-split-GAL4* driver lines that turn on prior to 36 hAPF, we quantified the fluorescence of V5-tagged Brp over metamorphosis (Supplemental Figure 11). We found Brp already present in LC4 at 36 hAPF, as supported by the RNAseq data, and that Brp expression increased until 60 hAPF. Unexpectedly, we witnessed a delay in the appearance and peak expression of Brp in LPLC2, similar to the staggered arrival times of LC4 and LPLC2 onto GF dendrites. It is possible that the assembly of synaptic machinery is delayed in LPLC2 to accommodate its arrival time. However, because V5-tagged Brp expression is dependent not only on the native *brp* promoter but also limited by when the VPN driver line turns on, the differences in Brp appearance could also be due to temporal differences in the driver lines. These data suggest presynaptic machinery is already present during initial partner matching between VPN and GF and increases as contacts are refined and stabilized.

We next investigated whether Brp accumulating at presynaptic terminals in VPNs was directly opposed to postsynaptic machinery, as an indicator of functional pre/postsynaptic sites. To label presynaptic Brp in VPN, we established a new transgenic line that expresses Brp-Short tagged with GFP under the control of the lexAop promoter (*lexAop-Brp-Short-GFP)*. Brp-Short is a truncated, non-functional Brp protein that localizes to sites of endogenous full-length Brp without disrupting morphology or function^47^ and has been used to map synaptic organization in the *Drosophila* CNS^48,49^. To label postsynaptic machinery in the GF, we targeted discs large 1 (*dlg1*), the fly PSD-95 ortholog^50^ using *dlg1[4K]*, a conditional tagging strategy that enables cell-type specific (*UAS-FLP*) V5-tagging of endogenous DLG1^49^. Combining these tools with our GF and VPN driver lines, we achieved co-expression of LC4-specific Brp-Short-GFP, and GF-specific DLG1-V5 and investigated protein expression patterns at distinct developmental stages. We observed faint, diffuse DLG1-V5 expression 36 hAPF (Figure 3C,D), around the time when initial GF and VPN contacts are observed (Figure 2F). However, significant DLG1-V5 and Brp-Short colocalization was not observed until 48 hAPF, although it remained only a small fraction of what was witnessed in the adult (Figure 3C,D). Our data suggest that although pre- and postsynaptic proteins are present at the initial stages of partner matching, it is not until around 48 hAPF that they begin to assemble functional synaptic connections.

### GF exhibits stimulus-independent neural activity during development

Our Brp-Short / DLG1-V5 dual labeling experiments suggest functional pre- and postsynaptic sites are present around 48 hAPF. Interestingly, within fly optic lobe neurons, sporadic and infrequent neural activity is first witnessed, through Ca^2+^ imaging, around 45 hAPF^51^. Developmental activity within DNs has not been investigated, so we set out to determine when activity first initializes within the GF and characterize GF activity patterns over development. We developed an *ex-plant* pupal electrophysiology preparation for high resolution recordings of the GF membrane potential over time (Figure 3E). Briefly, the entire pupal CNS was dissected and mounted onto a coverslip, which was then attached to a customized holder that enabled us to perfuse oxygenated extracellular saline during recordings. Using our preparation, we recorded from the GF for approximately one hour in current-clamp mode at distinct developmental time periods. At 45 hAPF, we witnessed sporadic, infrequent depolarizing events (Figure 3F and Supplemental Figure 12), aligning with the emergence of activity in the optic lobes^51^ and the initial opposition of Brp/DLG puncta (Figure 3C,D). The number of depolarizing events increased exponentially as development progressed (Figure 3F,G, Supplemental Figure 12), mirroring the increased expression of cholinergic synaptic machinery within the scRNA-seq data (Figure 3B, Supplemental Figures 9,10). As expected with an increase in the number of depolarizing events, the interval between events decreased (inter-event frequency increased) as a function of age (Figure 3I, Supplemental Figure 12).

Our recordings enabled us to observe hyperpolarizing events (Figure 3H, arrow) that occasionally proceeded large depolarizing events (Figure 3H, arrowheads), and small amplitude events that would not be resolvable with Ca^2+^ imaging. We also observed a broad distribution in event frequency (Figure 3G,I) instead of one dominant frequency from distinct alternating phases between silence and activity as seen in optic lobe or whole brain Ca^2+^ imaging from 55 - 65 hAPF^51,52^. It is possible our *ex-plant* preparation may alter activity patterns from those observed *in-vivo*. Alternatively, our recordings report activity not resolvable in Ca^2+^ imaging, and central brain neurons like the GF may display broader patterns as they pool input across many diverse cell types. Altogether, our data (summarized in Figure 3J) suggest initial GF partner matching precedes synaptogenesis. GF synapses become functional around 48 hAPF, with an upregulation of gap junction proteins and the appearance of apposed pre and postsynaptic machinery suggesting electrical (predominant) and chemical (minor) synapses contribute to the underlying activity witnessed at this stage. In the later stages of development, the frequency of synaptic events increase as gap junction proteins are downregulated and cholinergic presynaptic machinery is upregulated to enhance and stabilize synapses with intended synaptic partners while refining unintended contacts.

### LC4 ablation results in an increase of GF contacts with the LPLC2 glomerulus

After establishing our timecourse of GF and VPN interactions, we next investigated potential mechanisms that regulate VPN targeting and localization onto GF dendrites. Our data suggest synaptic activity does not contribute to the initial stages of VPN to GF partner matching. However, activity could be necessary for maintenance of the medial-lateral division of LC4 and LPLC2 inputs on GF dendrites, as neuronal activity can be crucial for proper refinement^53-57^. To test this, we attempted to silence LC4 during development by expressing the inwardly rectifying potassium channel Kir2.^17,58^ using *LC4_4-split-GAL4*. However, we found early expression of *Kir2.1* resulted in a significant loss of LC4 (Figure 4A; Supplemental Figure 13A-D). Expression of *Kir2.1* with a driver line used in previous silencing experiments^6^ that turns on later in development (*LC4_1-split-GAL4)* did not cause a loss of LC4 (Supplemental Figure 13A-D). As our *LC4_4-split-GAL4* driver line turns on ∼18 hAPF, prior to initial LC4 to GF contact, these data suggest that overexpression of Kir2.1 early in development is detrimental to LC4 survival, potentially due to the inability to compensate for disruptions in ionic homeostasis or the direct induction of apoptosis^59-61^. Co-expression of an apoptosis inhibitor p35^62^ with Kir2.1 using our *LC4_4-split-GAL4* driver line, however did not prevent cell death (Supplemental Figure 13E,F), potentially due to redundancies in apoptosis pathways or the relative timing of expression.

**Figure 4.**
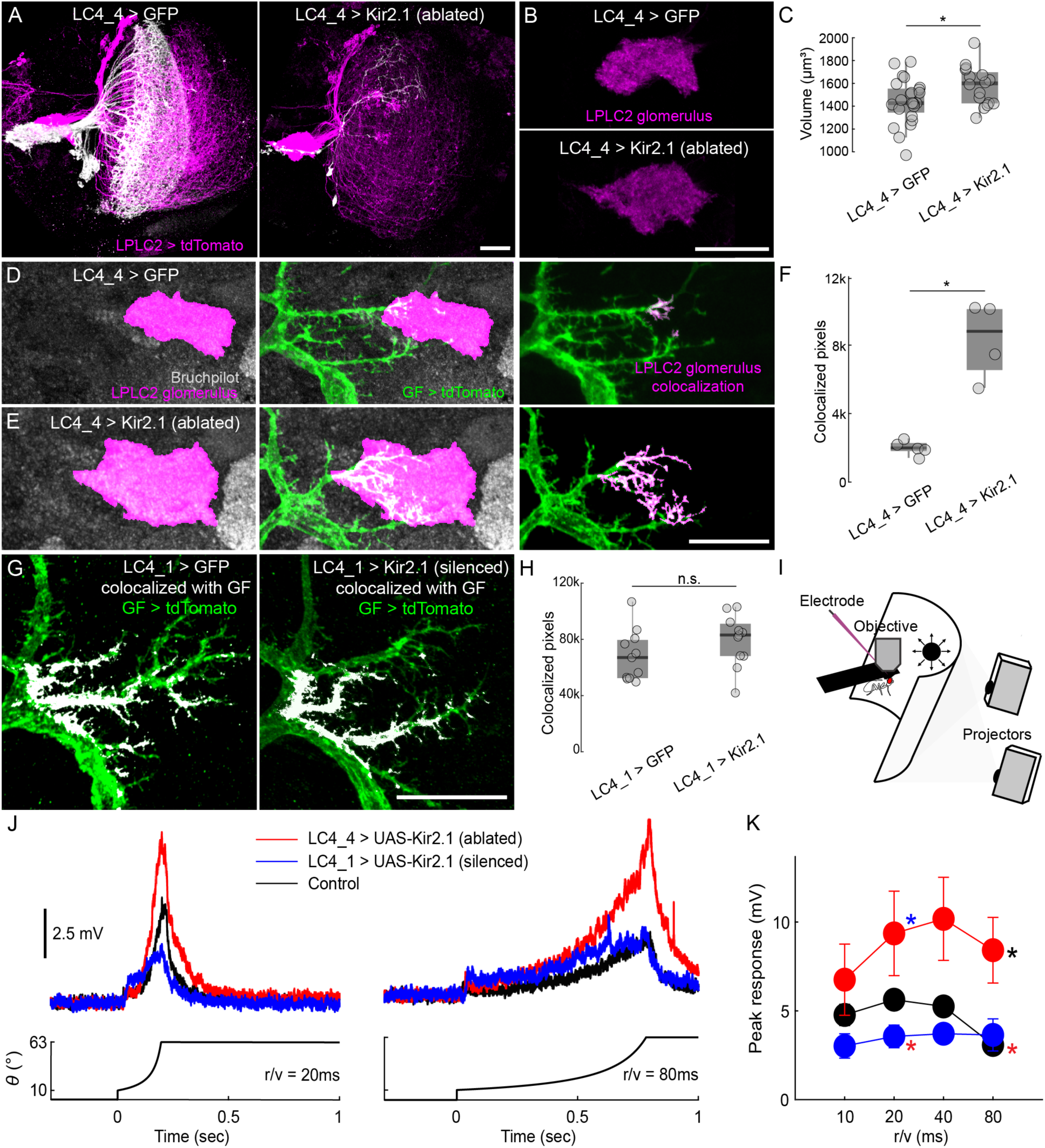
Developmental ablation of LC4 with Kir2.1 alters the morphology of the LPLC2 glomerulus and increases LPLC2 contacts and functional drive onto GF. **(A)** Maximum intensity projections of LPLC2 expressing tdTomato (magenta) and LC4 (white) expressing GFP (left) or Kir2.1 (right) using LC4_4, a driver line that turns on early in development. Scale bar, 20μm. **(B)** Maximum intensity projections of a substack of the LPLC2 glomerulus (magenta) in a fly where LC4 express GFP (top) or are ablated through Kir2.1expression (bottom). Scale bar, 20μm. **(C)** Quantification of LPLC2 glomerulus volume in (B). Two-sample *t-*test, p = .0273, N = 16-24 hemibrains from 8-13 flies. **(D)** Maximum intensity projections of Brp (NC82, gray) with the LPLC2 glomerulus highlighted (magenta) in a fly where GFP was expressed early in LC4 (left). Maximum intensity projections of GF dendrites (tdTomato, green) extending into the LPLC2 glomerulus (middle). Maximum intensity projections of colocalized pixels (magenta) between GF and the LPLC2 glomerulus superimposed onto the GF (right). **(E)** Maximum intensity projections of Brp (gray) with the LPLC2 glomerulus highlighted (magenta) in a fly where Kir2.1 was expressed early to ablate LC4 (left). Maximum intensity projections of GF dendrites (tdTomato, green) extending into the LPLC2 glomerulus (middle). Maximum intensity projections of colocalized pixels (magenta) between GF and the LPLC2 glomerulus superimposed onto the GF (right). Scale bar, 20μm. **(F)** Quantification of colocalization between GF and the LPLC2 glomerulus from (D,E). Unpaired Mann-Whitney U test, p = .0159, N ≥ 4 hemibrains from ≥ 4 flies. **(G)** Maximum intensity projections of GF expressing smGFP (green) with colocalized pixels (white) between LC4 expressing GFP (left) or silenced by Kir2.1 (right) using an LC4-split-Gal4 driver that turns on late during development (*LC4_1-split-GAL4*). Scale bar, 20μm. **(H)** Quantification of colocalization in (G). Unpaired Mann-Whitney U test, p = .1486, N ≥ 11 hemibrains from ≥ 6 flies. **(I)** Schematic representing *in-vivo* electrophysiology setup for head-fixed adult flies. Visual stimuli (looms) were presented ipsilateral to the side of the recording via projection onto a screen positioned in front of the fly. **(J)** Average GF responses to select looming stimuli presentations of different radius to speed ratios (r/v). **(K)** Quantification of peak amplitude responses to looming stimuli presentations in (J). Unpaired Kruskal-Wallis test (r/v = 10ms, p = .1105; r/v = 20ms, p = .0443; r/v = 40ms, p = .0556; r/v = 80ms, p = .0385), Tukey-Kramer multiple comparison test post hoc, * = p < .05, N = 6-8 flies.

With our finding we could use Kir2.1 as a tool to ablate LC4, we reframed our question to examine how the physical loss of LC4 alters LPLC2 morphology and targeting. Given that LC4 contacts GF dendrites ∼24 hours prior to LPLC2, we wondered if LC4 physically restricts LPLC2 from extending to medial regions of the GF optic glomeruli dendrites. We expressed tdTomato in LPLC2 using a LexA driver line (*LPLC2-LexA)* while simultaneously driving myrGFP or Kir2.1 with *LC4_4-split-GAL4*. In adult flies where Kir2.1 expression ablated the majority of LC4, the LPLC2 axon bundle extended into areas where the LC4 glomerulus would be expected and we witnessed a significant increase the LPLC2 glomerulus volume (Figure 4B,C). Due to the witnessed expansion of the LPLC2 glomerulus, we next investigated whether the loss of LC4 increased the territory GF dendrites occupied within the LPLC2 glomerulus. We expressed Kir2.1 with *LC4_4-split-GAL4* to achieve LC4 cell death, while simultaneously expressing tdTomato in GF using *GF_1-LexA.* We identified the LPLC2 glomerulus using a neuropil label (Brp) and again found that LC4 cell loss resulted in an LPLC2 glomerulus with altered morphology as compared to control flies (Figure 4D,E). We then quantified the overlap between GF dendrites and the LPLC2 glomerulus (Figure 4F). We found that LC4 ablation more than doubled the amount of colocalization between GF dendrites and the LPLC2 glomerulus, with the glomerulus expanding onto more medial regions of the GF dendrites as compared to controls (Figure 4D-F). In summary, our data demonstrate LC4 ablation results in altered LPLC2 axonal morphology and increased GF dendritic arborizations within the LPLC2 glomerulus, suggesting the early arrival and physical presence of LC4 may impede LPLC2 from contacting more medial regions of the GF.

To revisit our original question as to whether activity influences GF and VPN connectivity, we expressed Kir2.1 in LC4 using our late, *LC4_1-split-GAL4* driver line, and tdTomato in GF using our *GF_1-LexA* driver. The *LC4_1-split-GAL4* driver line should be effective at silencing LC4 as it expresses Kir2.1 prior to the onset of GF activity, as witnessed here (Figure 3F,G, Supplemental Figure 12), and Ca^2+^ activity, as observed in the fly’s visual system^51^. We found no significant difference in the density or localization of contacts between GF and LC4 whether we expressed Kir2.1 or myrGFP in LC4 (Figure 4G,H). These data suggest LC4 localization along GF dendrites is activity independent, and the early arrival and physical presence of LC4 axons restricts LPLC2 targeting to the lateral regions of GF dendrites.

### Functional compensation in the GF circuit occurs after LC4 ablation

Our anatomical data suggest the loss of LC4, but not silencing of its activity, during development results in a reconfiguration of contacts between LPLC2 and GF. We next investigated whether the apparent change in connectivity had functional consequences, affecting GF’s encoding of ethologically relevant visual stimuli. GF are tuned to looming stimuli – the 2D projections of an object approaching on a direct collision course. LC4 provides to the GF information about the angular speed while LPLC2 provides information about the angular size of a looming object^6,7^. We recorded GF responses in tethered, behaving flies using whole-cell patch clamp electrophysiology^63^ (Figure 4I). We displayed looming stimuli across different radius to speed (*r*/*v*) ratios where the contributions of LC4 and LPLC2 to the GF response have been previously established^6,7^. In control animals, LPLC2 contributions are maintained across stimuli, as the range in stimulus size does not change, while LC4 contributions increase as stimuli become more abrupt^6^.

We found, as reported previously^6^, that silencing LC4 by expressing Kir2.1 using the *LC4_1-split-GAL4* driver line reduced the GF response to looming stimuli as stimuli became more abrupt (Figure 4I-K). To further verify Kir2.1 silencing was effective, we expressed Kir2.1 in lamina monopolar cells 1 and 2 (L1-L2), the early-stage inputs to motion vision processing^64,65^ which ameliorated GF responses, as reported previously^6^ (Supplemental Figure 14). However, in contrast to what we witnessed with silencing, we found the ablation of the majority of the LC4 population, by expressing Kir2.1 with our *LC4_4-split-GAL4* driver line, resulted in an enhanced GF response to looming stimuli (Figure 4I-K), aligned with the observed increase in GF dendrites occupying the LPLC2 glomerulus (Figure 4D-F). Our data support that the developmental reorganization of LPLC2 and GF following LC4 ablation is functionally significant and leads to an over-compensation in the GF looming response.

## DISCUSSION

Here, our investigation into the interactions of GF dendrites and VPN axons support a developmental program where GF and partner VPNs make initial contact in precise, stereotyped regions that are maintained into eclosion through competitive, physical interactions. Ablation of one major VPN partner (LC4) results in territory expansion of another VPN (LPLC2) that confers compensatory functional changes within the GF circuit. After initial VPN territories are established, GF dendrites continue to arborize and increase contacts with VPNs, while avoiding contacts with non-synaptic neighbors. This developmental stage coincides with an upregulation of gap junctions, the opposition of pre- and postsynaptic proteins, and the onset of developmental activity in the GF. This outgrowth is followed by a period of stabilization/refinement that coincides with an upregulation of cholinergic synaptic machinery and an increase in the frequency of developmental activity in the GF. This developmental time course is summarized in Figure 3A,J. Our data establish the GF escape circuit as a sophisticated developmental model that can be used to study mechanisms establishing integration of sensory inputs within a sensorimotor circuit, the role neural activity plays in shaping circuit connectivity and refinement, and the relationship between expressed genes and circuit development and function.

The development of the *Drosophila* visual system proceeds in a series of steps which likely serve to reduce the complexity of wiring paradigms from neurons of the same or neighboring cell-types, highlighting the importance of timing^66^. We found staggered interactions of the GF with VPN partners, where LC4 contacts the GF approximately 24-36 hours prior to LPLC2. This staggered arrival of VPN axons could reduce the complexity of decisions made by the GF during partner matching, as is also seen in the olfactory glomeruli in an *ex-plant* preparation where axons of pioneer olfactory receptor neurons (ORNs) terminate in posterior regions, and ORNs arriving later terminate in anterior regions^67^. From 36-72 hAPF, we observe an increase in GF dendritic complexity and extension and increase in contacts with VPN partners, coordinated with an upregulation of genes involved in the SNARE complex (Figure 3A,B). This period of precise targeting and outgrowth likely reflects a robust partner matching program, potentially through ligand-receptor or attractive/repulsive cues^68^. Our confocal data provide high-resolution snapshots of membranes at distinct periods over metamorphosis, but as metamorphosis is a dynamic process, future work could incorporate time-lapsed imaging to investigate transient interactions that may have been missed.

Our GF and VPN colocalization data show a developmental progression that begins with partner matching between VPN and GF in stereotyped regions, an increase of contacts with synaptic partners, proceeded by refinement as neurons assume their adult morphology. Comparison with the time course of scRNA-seq data provides insight into genes that may play a role in these processes. We find gap junction coupling may serve a role in partner matching, as *shakB* expression is high at this time (Supplemental Figure 9)^22^. Our data also support a transition from predominantly electrical to chemical synapses as witnessed in other species^69,70^ at the onset of refinement. *shakB* is downregulated while cholinergic synaptic machinery *ChAT* and *VAChT* (Figure 3B and Supplemental Figures 9,10) are upregulated. Our model system is well poised to investigate the role of electrical and chemical signaling, and their supporting genes, in circuit development and function.

We provide the first electrophysiological recordings of developmental neural activity in pupal neurons, a phenomenon that has been documented in developing vertebrate systems, and recently proposed within the fly with Ca^2+^ imaging^53-57,69,71-75^. Our data demonstrate activity in the GF emerges as early as 45 hAPF, and increases in frequency as a function of time. While scRNA-seq data and our Brp-Short and DLG1-V5 protein expression data suggests this activity is driven through functional electrical and chemical synapses, changes in GF intrinsic properties may also contribute to witnessed changes in frequency. For example, recordings from the superior olivary nucleus in the avian auditory brainstem over embryonic development to hatching show neuronal excitability increases due to changes in K^+^ and Na^+^ ion channel conductance^76^, therefore further investigation into what drives these changes in GF activity is warranted.

To record spontaneous activity in pupal GF neurons we developed an *ex-plant* system that differs from established *ex-vivo* systems^77-79^ in that we are not attempting to culture our ex-plant long term and replicate *in-vivo* conditions through the addition of ecdysone to aid neuronal development. To our knowledge, developmental activity within these ex-vivo systems has yet to be reported. We find in our ex-plant noticeable similarities to the recent discovery of *in-vivo* activity patterns in the developing fly nervous system^51^. We first observe GF stimulus independent activity (stimulus independent because all sensory organs are no longer connected) as infrequent events around 45 hAPF, similar to sparse Ca^2+^ activity witnessed in the fly optic lobes at this same time^51^. As development progresses, the frequency of depolarizing events in the GF increases, similar to what has been reported *in-vivo.* However, we find no discernible phases of activity and silence as observed *in-vivo* via calcium imaging around 55-65 hAPF, classified as the periodic stage of patterned, stimulus independent neural activity (PSINA). We instead witness a progression into what resembles the later turbulent phase of PSINA that occurs around 70 hAPF to eclosion^51^. As in-vivo activity in individual developing DNs or central brain neurons has as of yet to be reported, our data could represent actual in-vivo activity patterns from neurons in the central brain where multiple inputs converge. Alternatively, even if our removal of the CNS disrupts activity patterns observed in-vivo, our ex-plant could provide a highly accessible model system to uncover the underlying mechanisms for how particular activity patterns arise.

The location of a synapse on a dendrite can impact its overall effect on a neuron, and establish how it contributes to neural computations^80-82^. We find the location of synapses of each VPN cell type to be highly stereotyped, suggesting location may impact computation, although this has yet to be directly investigated. Our contact data suggest that targeting of LC4 and LPLC2 to their respective regions is established upon initial contact instead of refinement in the later stages of development. This is noticeably different from ORN axonal targeting to olfactory glomeruli, where axons target many neighboring glomeruli and are eventually refined to a specific glomerulus^67,83,84^. It is possible specific protein interactions establish LC4 and LPLC2 target specificity^68,85-87^, similar to how basket interneurons target the axon initial segment of Purkinje cells in the vertebrate cerebellum via localized cell adhesion molecules and adaptor proteins^88-91^. Our data, however, support that axon arrival times also play a role, with LC4 first contacting GF dendrites in medial regions, physically impeding LPLC2, and leaving LPLC2 segregated to the lateral regions. This physical barrier may explain why LPLC2 is able to extend into medial regions following LC4 ablation (Figure 4D-F). We do however find LPLC2 does not fully replace LC4 along the dendrites, suggesting segregation may arise from a combination of physical restraints from LC4 and potentially other neurons, in addition to molecular interactions. It does not appear that activity-based mechanisms influence the localization of LC4 and LPLC2 to specific regions because silencing activity in LC4 does not result in significant changes in LC4/GF contact density or localization (Figure 4G,H). In addition, elimination of the majority of LC4 neurons did not affect targeting of the remaining LC4 neurons to the GF dendrites in the expected regions.

We also report altered GF output to looming stimulus presentations (Figure 4J,K) following LC4 ablation as an example of developmental plasticity to preserve an evolutionary conserved escape behavior that is critical to the fly’s survival. Because we observe a significant increase in GF dendritic occupancy within the LPLC2 glomerulus, our prevailing hypothesis is that LPLC2 synaptic inputs to GF have increased. Alternatively, the 1-7 LC4 neurons that remain after ablation may also have increased synaptic input, however the consistency of the compensatory responses as looming stimulus parameters change (*r*/*v*) and the limited visual field coverage with just 1-7 LC4 neurons supports compensation through remaining LC4 is unlikely to underlie the enhanced GF responses.

While we here demonstrate stereotyped targeting of VPNs to distinct regions on GF dendrites, emerging work suggests an additional level of targeting may occur within individual VPNs. It was previously suggested, based on light microscopy data, that retinotopy is lost within the seemingly random terminations of VPN axons in most optic glomeruli, unlike in vertebrate systems where established retinotopy in the retina is maintained in projections to the lateral geniculate nucleus and superior colliculus/tectum^92-94^. However, recent evidence suggests VPNs preserve spatial information by biasing synaptic input to postsynaptic neurons relative to their receptive field^8,10,19^. This is seen in LC4 synaptic inputs to postsynaptic DNp02 and DNp11 neurons, where LC4 synaptic inputs to DNp02 increase along the posterior to anterior visual axis, and LC4 synaptic inputs to DNp11 increase along the anterior to posterior visual axis^10^. LC4 inputs to GF do not appear to bias synapse numbers based on their receptive field, but a bias of synaptic inputs from LPLC2 to GF exists along the ventral to dorsal axis^10^. As synaptic gradients appear to be utilized amongst many VPN neurons, our model system is well poised to investigate when and how these synaptic biases arise.

In summary, our data provides a detailed anatomical, transcriptomic, and functional description of GF and VPN development. Our model is unique in that we can observe multiple visual feature inputs competing for dendritic space, providing a complex sensorimotor model to the field that will be useful to determine the relationship between connectivity and sensorimotor integration. The GF also receives input from other brain regions outside of the optic glomeruli^95^, and it would be interesting to characterize the development of GF with respect to these other regions and further investigate how these inputs influence GF output. Finally, these VPN are only a few of the 20+ VPN that terminate in the optic glomeruli, and GF is one of hundreds of DNs, expanding the opportunity to uncover conserved, fundamental mechanisms for the wiring of sensorimotor circuits.

## METHODS

### Fly genotypes and rearing

Drosophila stocks (Table 1) and experimental crosses (Supplementary Table 1) were reared on a traditional molasses, cornmeal, and yeast diet (Archon Scientific), maintained at 25°C and 60% humidity on a 12-hour light/dark cycle, except for optogenetics experiments where dark reared flies were raised on 0.2 mM retinal food as larva and switched to 0.4 mM retinal food following eclosion. All experiments were performed on pupal or adult female flies 2-5 days post-eclosion. New split-GAL4 drivers lines SS02569 and SS02570 were generated using previously described methods^11^. The Janelia FlyLight Project Team contributed to split-GAL4 screening and stock construction.

**Table 1.**
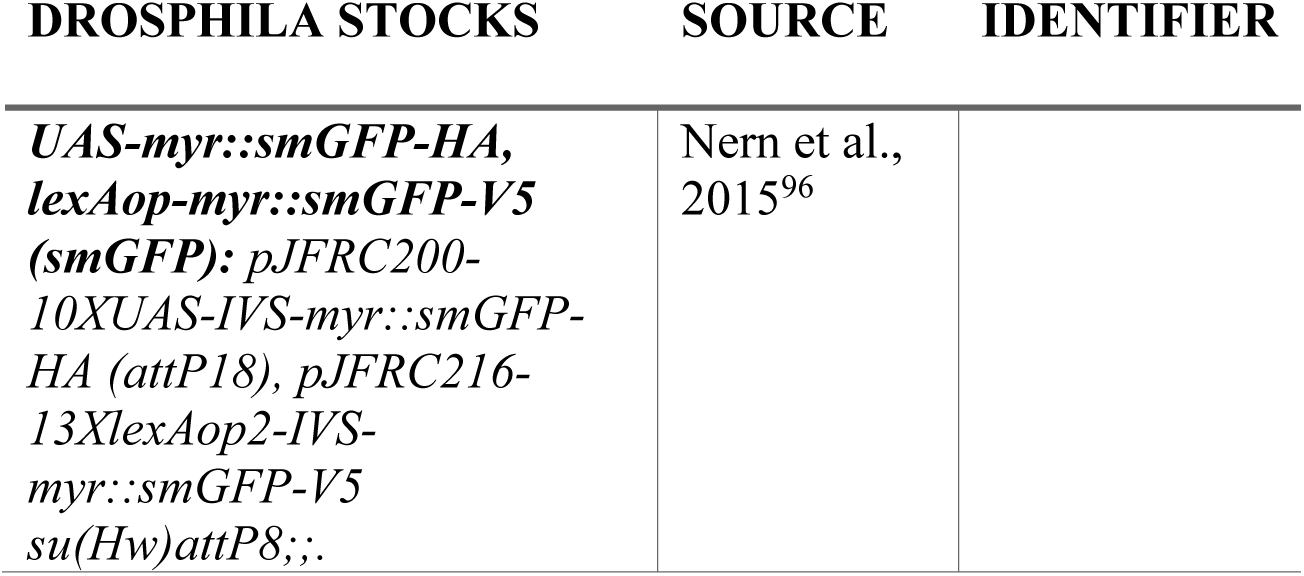
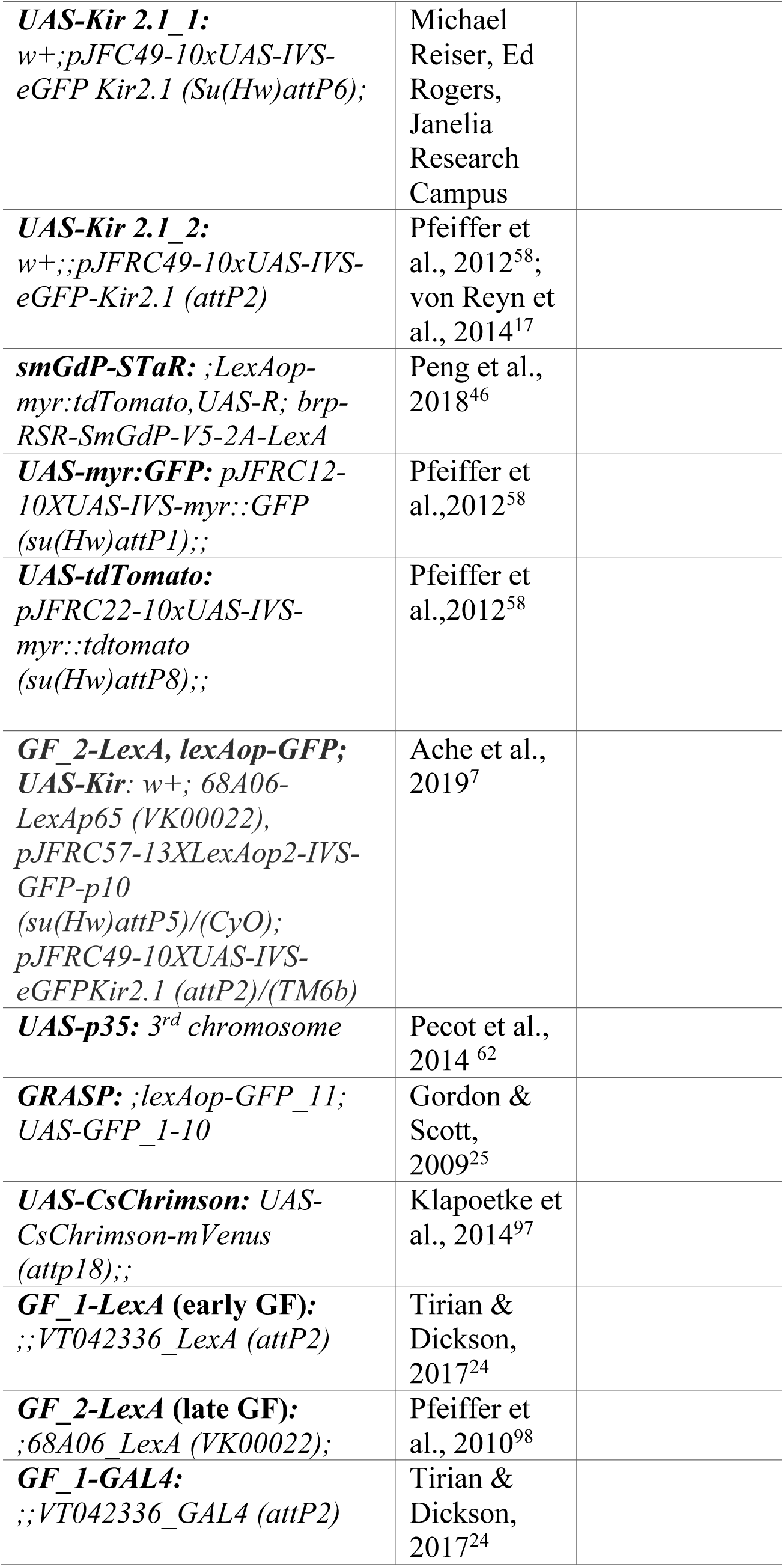
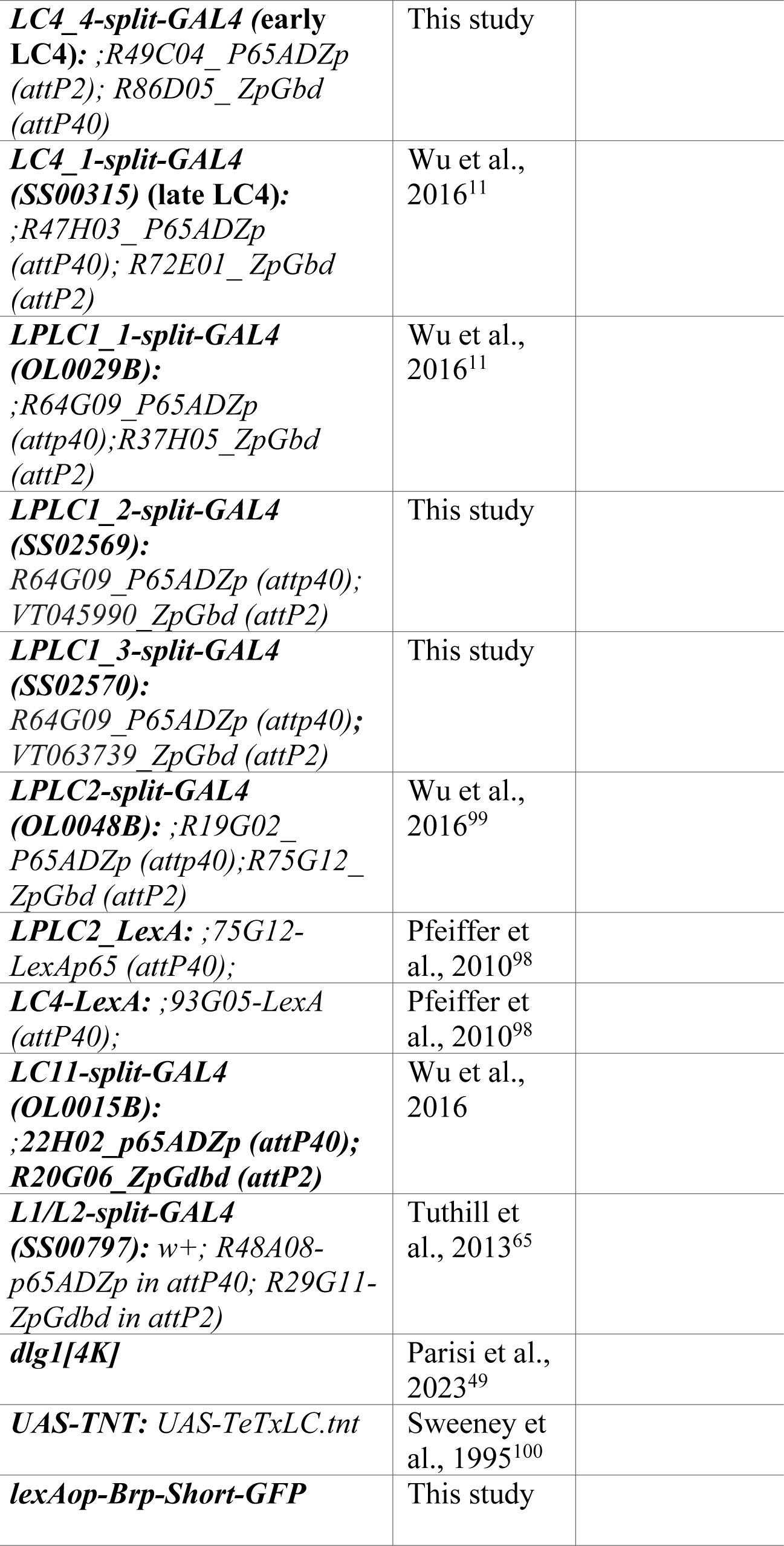
*Drosophila* Stocks.

### Developmental staging

Pupal staging across developmental time points has been previously described^99^. In brief, the sex of white pre-pupa was identified, and females were transferred to a separate petri dish, marked as 0 hours after pupa formation (hAPF), and reared for the appropriate amount of time at 25°C before dissection. Dissections were performed within a 2-hour window of a targeted pupal developmental stage. All pupal dissections were synchronized and processed through immunohistochemistry protocols for pixel intensity measurements of images.

### Immunohistochemistry

All dissections were performed in cold Schneider’s insect media (S2, Sigma Aldrich, #S01416) within a 15-minute window before solution exchange to avoid tissue degradation. Brains were then transferred to a 1% paraformaldehyde (20% PFA, Electron Microscopy Sciences, #15713) in S2 solution and fixed overnight at 4°C while rotating. Immunohistochemistry was performed as described previously^96^. Primary and secondary antibodies are listed in Table 2. Supplementary Table 1 lists antibodies used for each figure with their respective dilutions. Following immunostaining, brains mounted onto poly-L-lysine (Sigma Aldrich, #25988-63-0) coated coverslips were dehydrated in increasing alcohol concentrations (30, 50, 75, 95, 100, 100) for 5 minutes in each, followed by two 5-minute Xylene clearing steps (Fisher Scientific, #X5-500). Coverslips were mounted onto a prepared slide (75 x 25 x 1 mm) (Corning, #2948-75×25) with coverslip spacers (25 x 25 mm) (Corning, #2845-25) placed on each end of the slide to prevent brain compression. Brain mounted slides were left to dry for at least 48 hours prior to imaging.

**Table 2.**
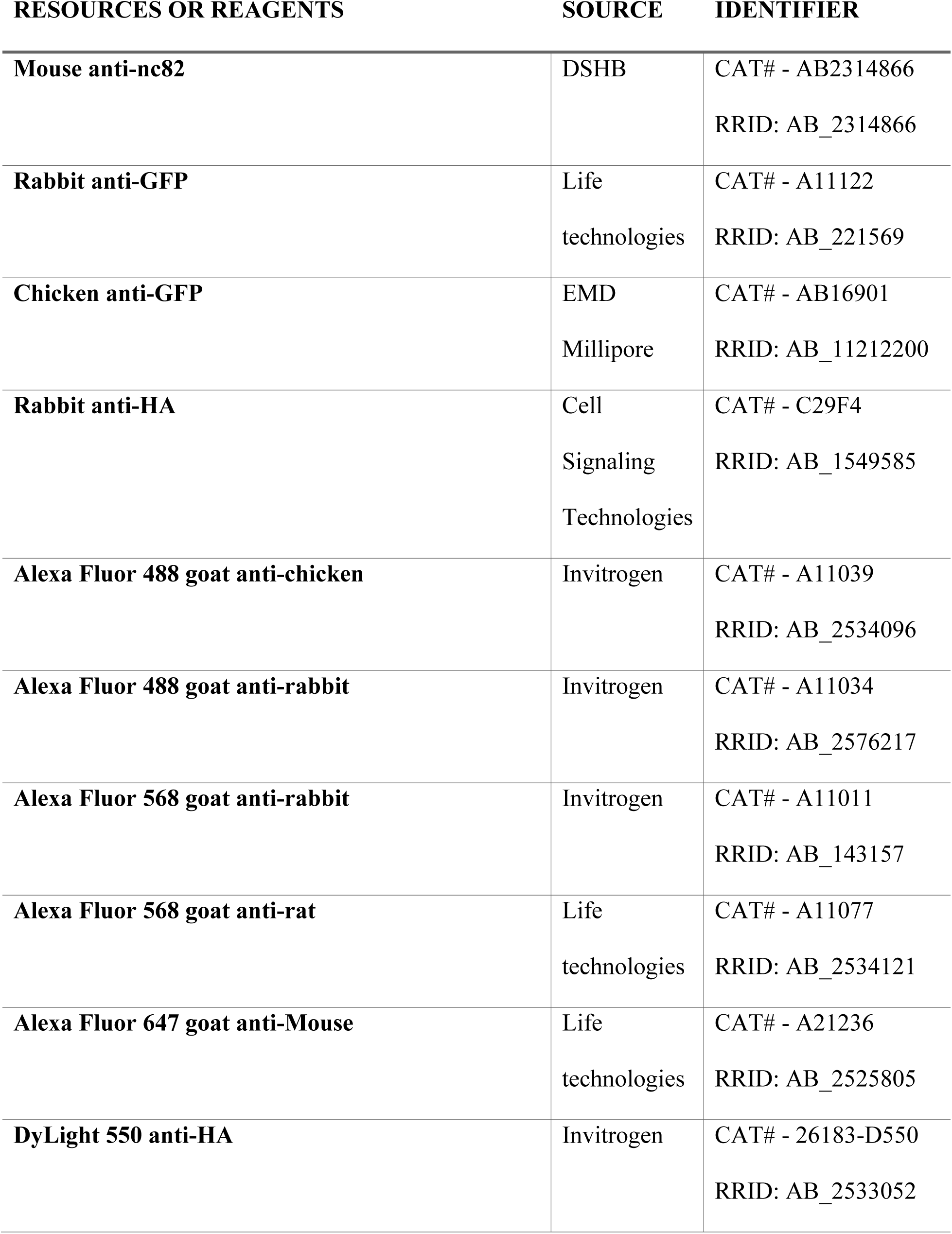
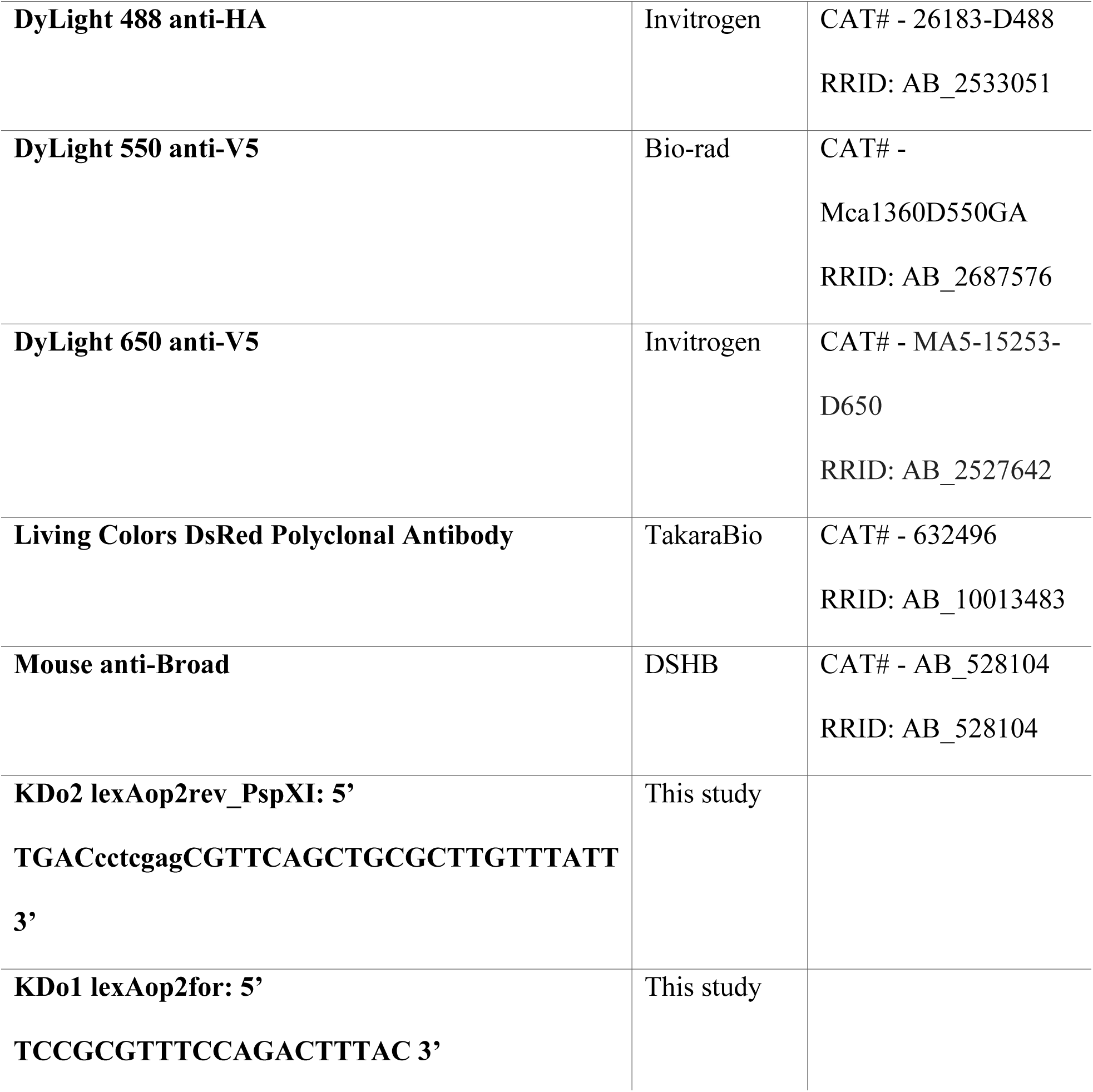
Resources and Reagents.

### Confocal Microscopy

Unless otherwise stated, all images were taken on an Olympus Fluoview 1000 confocal system. Images were taken with a 60x, 1.42 NA oil immersion objective to achieve a voxel size of .103µm x .103µm x .45µm. Imaging parameters were minimally adjusted between images to achieve an image that utilizes the full pixel intensity range without oversaturating pixels. This was necessary as driver lines in earlier pupal developmental periods showed lower levels of expression than in later pupal developmental stages, therefore imaging parameters were adjusted to optimize the membrane signal-to-noise ratio for each developmental stage that would allow for optimized mask generation used in image analysis. In analyses where pixel intensities were compared across developmental stages, all imaging parameters were kept consistent across all images. STaR images were taken on a Zeiss LSM 700 with a 63x, 1.4 NA oil immersion objective to achieve a voxel size of .06µm x .06µm x .44µm. Imaging parameters were kept consistent to allow for comparison across all samples. Images for LPLC2 glomerulus volume quantification were taken on a Zeiss LSM 700 with a 63x, 1.4 NA oil immersion objective with a magnification of 0.50 to achieve a voxel size of .0992µm x .0992µm x .3946µm.

### Electron Microscopy

The publicly available electron microscopy hemibrain dataset (version 1.2.1)^21^ was used in this paper. NeuPrint^23^ was used to create renderings and connectivity diagrams.

### Image analyses

A region of interest (ROI) was drawn around the GF optic glomeruli dendrites to quantify dendritic complexity and length. To quantify dendritic complexity, all pixels in this ROI were summed. The Euclidean distance was measured from the beginning (most medial aspect) of the GF optic glomeruli dendrites to the tip to calculate dendritic length.

To quantify membrane colocalization between GF and VPN neurons across development, intensity-based thresholding was first used to generate a binary mask of each neuron membrane. Using a custom GUI written in MATLAB, threshold values were manually selected to include processes of each cell-type while excluding background and regions of the neuron that were out of focus in each z-plane. Generated binary masks were inspected to make sure each mask was representative of the imaged neuron membrane channel. In certain cases, the set threshold did not include very fine neurites that were difficult to discriminate from background. Lowering threshold values to capture these processes in the mask would result in background being included into the mask as well, therefore generated masks may exclude some of the finer dendritic processes with low SNR. Using these masks, colocalized pixels were collected plane-by-plane across the entire image stack using Boolean operators between GF and VPN masks.

This output matrix resulted in a z-stack where only GF and VPN membranes were colocalized. In some images, GF membrane labeling had a low SNR and prevented accurate mask generation and was therefore excluded from analyses. In some cases, non-GF and non-VPN cell-types were labeled with the driver lines, therefore these data were excluded from analyses. To increase rigor, a second method to quantify GF to VPN membrane colocalization was used. Confocal stacks of labeled neuronal membranes were 3D rendered in Imaris using the Surfaces function. Thresholds were manually applied to generate 3D masks of neuronal membranes so that the rendered 3D image was representative of the imaged membrane. Similar to MATLAB thresholding, in certain cases the set threshold did not include fine neurites as they were difficult to discriminate from the background using automated algorithms. Following initial 3D membrane renderings, renderings were inspected and regions where faint processes were still visible but not detected in the thresholding pipeline were manually filled in using the ‘Magic Wand’ function. Once 3D rendering was complete, areas of colocalization were identified using the ‘Surface-Surface contact area’ XTension, and the volume of this output was quantified.

To determine the density of VPN contacts along the GF optic glomeruli dendrite, a z-projection of the GF was manually aligned and rotated using anatomical landmarks consistently observed to align the lateral dendrite along the medial-lateral axis (x-axis). The same rotation and alignment were applied to the appropriate membrane colocalization matrix. An ROI was then drawn around the GF optic glomeruli dendrites, with anatomical landmarks consistently observed used to denote the beginning and end of the optic glomeruli dendrite. To account for variation in the optic glomeruli dendrite extension across images, we normalized the x-axis to the length of the drawn ROI (i.e., GF lateral dendrite). To determine where VPN contacts were localized along the dorsal-ventral axis (y-axis), the same images were used, but normalized the y-axis to the length of the drawn ROI. Total colocalized pixels were summed along each column or row, respectively, and the pixel density for each column or row was averaged across brains in each condition, then plotted along the normalized axis.

To quantify the average pixel intensity of VPN-specific Bruchpilot (Brp) puncta across developmental timepoints, VPN masks were generated from the membrane channel using the same pipeline used for the GF and VPN membrane colocalization. These binary masks were then multiplied to the Brp-puncta channel to gather raw pixel intensities for Brp-V5 puncta localized to the VPN membrane. The total intensity sum of the glomerulus was divided by the total number of membrane localized pixels to calculate the average pixel intensity.

To isolate Brp pixels that colocalized with DLG, the DLG channel was first thresholded using FIJI’s default auto-thresholding function, binarized in MATLAB, and then multiplied to the Brp channel. An ROI mask was then used to restrict Brp analysis to the VPN glomerulus of interest. The total number of Brp positive pixels was then calculated and the overall volume of Brp-DLG colocalization computed by multiplying the total pixel count by the image voxel size.

To quantify LPLC2 glomerulus volume, the LPLC2-membrane channel and Brp channel were thresholded in Fiji and binarized using MATLAB as described above. The two channels were then multiplied where pixels containing both membrane label and Brp were considered part of the glomerulus. Glomerulus volume was determined by multiplying the total glomerulus pixel count by image voxel size.

For analyses where dendrite complexity and extension were quantified (Figure 2B,C), a median filter was used to remove background noise. For all other images, no pre-processing was performed, and only the brightness and contrast were adjusted to highlight neuronal processes when preparing images for figure generation. For all data sets using the GF-LexA driver line, any images that had non-GF cell types within our ROI and low or no GF expression because of driver line stochasticity were excluded from analyses.

### Creation of the LexAop2-Brp-Short-GFP-HSV Transgenic Line

To create a transgenic line expressing Brp-Short-GFP under control of the lexAop promoter (lexAop-Brp-Short-GFP), we used the Gateway cloning system (Thermo Fisher Scientific, cat. No. K202020) via an existing plasmid containing the UAS-Brp-Short sequence^101^ followed by a Gateway cassette. We excised the UAS sequence using dual HindIII and PspXI restriction digests and replaced the promoter with a lexAoperon sequence, flanked by HindIII and PspXI restriction sites, that was first PCR amplified using custom primers (see Table 2) from a plasmid containing lexAop2 (lexAop2-myr-4xSNAPf, RRID: Addgene 87638) and then restriction digested using HindIII and PspXI (New England BioLabs, Ipswich, MA) to create compatible sticky ends. Following ligation and confirmation of the appropriate promoter insertion by sequencing, we replaced the Gateway cassette with the GFP-HSV tag from an Entry vector via Gateway LR recombination reaction (Thermo Fisher Scientific, cat. no. 11791019). Plasmid identity and the presence of all components was verified by sequencing (GeneWiz, South Plainfield, NJ). Transgenic lines of the resultant plasmid inserted into the φC31 site at attP2 (Bloomington Drosophila Stock Center, RRID: 8622) located at 68A4 on the 3^rd^ chromosome were then produced using standard methods (BestGene, Inc., Chino Hills, CA). Subsequent lines were verified by genomic sequencing and a single line chosen for experiments.

### Kir2.1 cell death and GF dendrite localization

To quantify LC4 cell death following early expression of Kir2.1, immunohistochemistry was performed against a GFP conjugated to Kir2.1 or GFP (controls), and Brp. Following imaging, LC4 cell bodies were manually counted using the GFP channel. To quantify GF dendrite density within the LPLC2 glomerulus following LC4 cell death, the Brp channel was used to visualize the optic glomeruli active zones. Axons that make up each individual glomerulus reliably terminate in the same region of the central brain, allowing for consistent identification of the LPLC2 glomerulus.

### scRNA-seq data analysis

To quantify changes in mRNA expression over development for our cells of interest, a recently published scRNA-seq dataset was used^22^. For each developmental stage for each population, an unpaired non-parametric Kruskal-Wallis test by ranks was performed, followed by a Dunn-Bonferroni multiple comparisons test for significant groups.

### Electrophysiology

For adult whole-cell electrophysiology, female flies were head fixed to recording plates via UV glue, antenna were UV-glued, and the front legs were removed at the level of the femur as described previously^6,63^. GFP positive GF soma were accessed for recordings by removing the cuticle and overlying trachea, and then removing the perineural sheath by local application of collagenase (0.5% in extracellular saline). Brains were perfused with standard extracellular saline (103 mM NaCl, 3 mM KCl, 5 mM *N*-Tris (hydroxymethyl)methyl-2-aminoethane-sulfonic acid, 8 mM trehalose, 10 mM glucose, 26 mM NaHCO3, 1 mM NaH2PO4, 1.5 mM Cacl2 and 4 mM MgCl2, pH 7.3, 270–275), bubbled with 95% O2/5% CO2 and held at 22°C. Recording electrodes (3.5-6.2 MΩ) were filled with intracellular saline (140 mM potassium aspartate, 10 mM HEPES, 1 mM EGTA, 4 mM MgATP, 0.5 mM Na3GTP, 1 mM KCl, 20 μM Alexa-568-hydazide-Na, 260-275 mOsm, pH 7.3). Recordings were acquired in whole-cell, current clamp mode, digitized at 20kHz, and low pass filtered at 10kHz. All data were collected using Wavesurfer, an open-source software (https://www.janelia.org/open-science/wavesurfer) running in MATLAB. Recordings were deemed acceptable if a high seal was attained prior to break through, the resting membrane potential was ≤ −55 mV, and the input resistance was > 50 MΩ. Current was not injected to hold the membrane potential at a particular resting level, and traces were not corrected for a 13mV liquid junction potential^102^.

All pupal recordings were staged in accordance with our staging protocol. Extracellular and intracellular reagents used were identical to the reagents used for adult recordings. Recordings were acquired in whole-cell, current clamp mode, digitized at 20kHz, and low pass filtered at 10kHz. All data were collected using Wavesurfer running in MATLAB. Recordings were deemed acceptable if recording electrodes (3.4 – 5.2 MΩ) attained a high seal (GΩ range) prior to break through, the resting membrane potential was below ™30mV and remained stable throughout the duration of the recording, and the input resistance ranged from 50 MΩ to 300 MΩ. Current was not injected to hold the membrane potential at a particular resting level, and traces were not corrected for a 13mV liquid junction potential^102^.

### Optogenetics

Light activation of VPN cell types expressing CsChrimson^97^ while recording from GF was performed by delivering light (635nm LED, Scientifica) through a 40x objective focused on a head fixed fly. Light pulses (5ms,1.7 μW/mm^2^, as measured in air at the working distance of the objective) were delivered 5 times at 30 second intervals.

### Visual Stimuli

Visual stimuli were projected on a cylindrical screen surrounding a head fixed fly during whole-cell electrophysiology following the protocol described previously^63,103^. A 4.5-inch diameter mylar cylindrical screen covered 180° in azimuth, and two DLP projectors (Texas Instruments Lightcrafter 4500) were used to minimize luminance attenuation at the end of the screen edge. The projections from the two projectors were calibrated on the cylindrical screen surface as described previously^103^ and the two projections overlapped 18° in azimuth at center of the screen and blended for uniform illumination. Generated looming stimuli based on the equation^104^ below and constant velocity expansion stimuli were displayed with 912 x 1140 resolution in 6bit grayscale at 240 Hz which is above the flicker fusion frequency of *Drosophila* (100 Hz^105^). Looming stimuli were generated by simulating a 2D projection of an object approaching at a constant velocity which mimics an approaching predator. The angular size (θ) of the stimulus subtended by the approaching object and was calculated over time (*t*) by the following equation^104^:

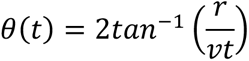

where *t*<0 before collision and *t*=0 at collision for an approaching object with a half size (r) and constant velocity (v). Four looming stimuli (r/v = 10, 20, 40, 80ms) were displayed, starting at 10°, expanding to 63° and then held for 1 second. Stimuli were presented once per trial, in a randomized order, every 30 seconds. For each fly, two trials of the entire set of stimuli were averaged.

### Data analysis and Statistics

No power analysis was performed prior to statistical analysis. Volume analysis in Figure 3 was performed by a researcher blinded to the genotypes. All data from confocal microscopy experiments were tested for normality using a Kolmogorov-Smirnov test or Anderson-Darling test and the appropriate parametric or non-parametric test was performed, as stated in the figure captions.

For boxplots, the dividing line in the box indicates the median, the boxes contain the interquartile range, and the whiskers indicate the extent of data points within an additional 1.5 × interquartile range.

For *in-vivo* electrophysiology analyses in adult recordings, all analyses were performed using custom MATLAB scripts. Recordings for each stimulus presentation were baseline subtracted by taking the average response one second prior to the stimulus onset. The magnitude of the GF expansion peak was measured after filtering each recording (Savitzky–Golay, fourth order polynomial, frame size is 1/10th the length of the stimulus). The normality of the data was assessed using a Kolmogorov-Smirnov test. If the data were found to not follow a normal distribution, the appropriate non-parametric test was selected. For non-parametric analyses, a Kruskal-Wallis test was performed, and Tukey-Kramer post hoc test was performed for significant groups.

For *ex-plant* pupal recordings, all analyses were performed using in-house MATLAB scripts. Potential 60Hz noise was filtered out using a band-stop filter, and thirty minutes of data were quantified. The baseline was determined after two rounds of computing the average signal envelope. Peaks were identified by capturing all depolarizations that were 3mV above baseline and separated by at least 100ms. Time intervals between events were transformed into instantaneous frequency for histogram plots.

## ACKNOWLEDGEMENTS

We thank the Janelia FlyLight project team for assistance with split-GAL4 generation. This study was supported in part by the National Science Foundation (grant no. IOS-1921065 to C.R.v.R.), the National Institutes of Health (NINDS R01NS110907 to T.J.M. and NINDS R01NS118562 to C.R.v.R.) and the Margaret Q. Landenberger Research Foundation (grant to C.R.v.R.).

## AUTHOR CONTRIBUTIONS

Conceptualization C.R.v.R.; Methodology, B.W.M and C.R.v.R.; Investigation, B.W.M., H.J., Y.Z.K., N.S., B.W.H. and C.R.v.R. Writing – Original Draft, B.W.M. and C.R.v.R.; Writing – Review & Editing, B.W.M., N.S., H.J., B.W.H., T.A.G., Y.Z.K., A.N., M.J.P, K.C.D, T.J.M., and C.R.v.R.; Funding Acquisition, C.R.v.R.; Resources, M.J.P, K.C.D, T.J.M., T.A.G., Y.Z.K. and A.N.; Supervision, C.R.v.R.

## DECLARATION OF INTERESTS

The authors declare no competing interests.

## Notes

### Competing Interest Statement

The authors have declared no competing interest.

### Summary of Updates

Updated funding information Revised Supplemental Figure 12

