## Supplemental Figures and Tables for "Axon arrival times and physical occupancy establish visual projection neuron integration on developing dendrites in the *Drosophila* optic glomeruli"

**SUPPLEMENTAL FIGURES AND TABLES (McFarland et al. 2024)**

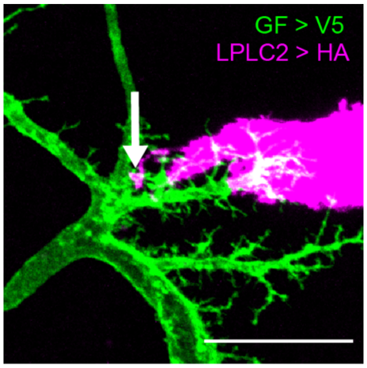

**Supplemental Figure 1. LPLC2 mistargeting onto the medial regions of GF dendrites**
Maximum intensity projection of the adult GF (green) with respect to LPLC2 (magenta). Arrow indicates a single LPLC2 axon extending into the medial region of the GF dendrite where LC4 predominantly resides. Scale bar, 20μm.

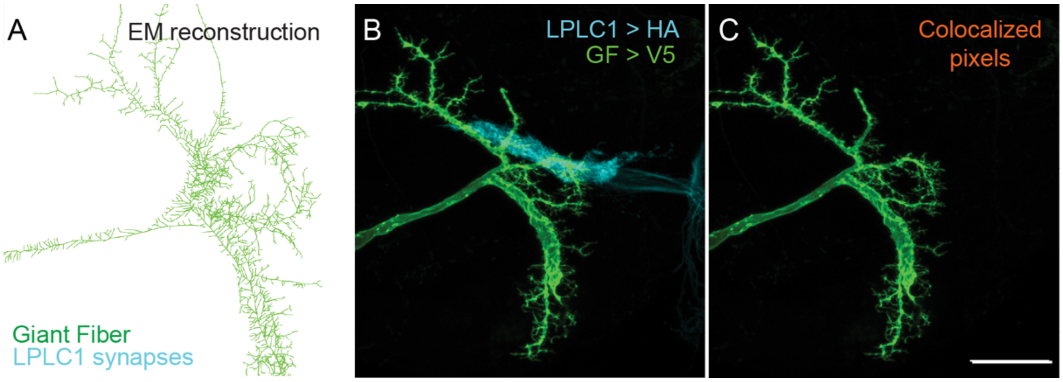

**Supplemental Figure 2. LPLC1 and GF are not synaptically coupled in adults**
**(A)** *Drosophila* hemibrain EM reconstruction of adult GF (green) with colored dots indicating input synapses from LPLC1 (cyan).
**(B)** Maximum intensity projection of dual labeled GF (green) and LPLC1 (cyan).
**(C)** Maximum intensity projection of colocalized pixels (orange) between GF and LPLC1 superimposed over GF. Scale bar, 20µm.

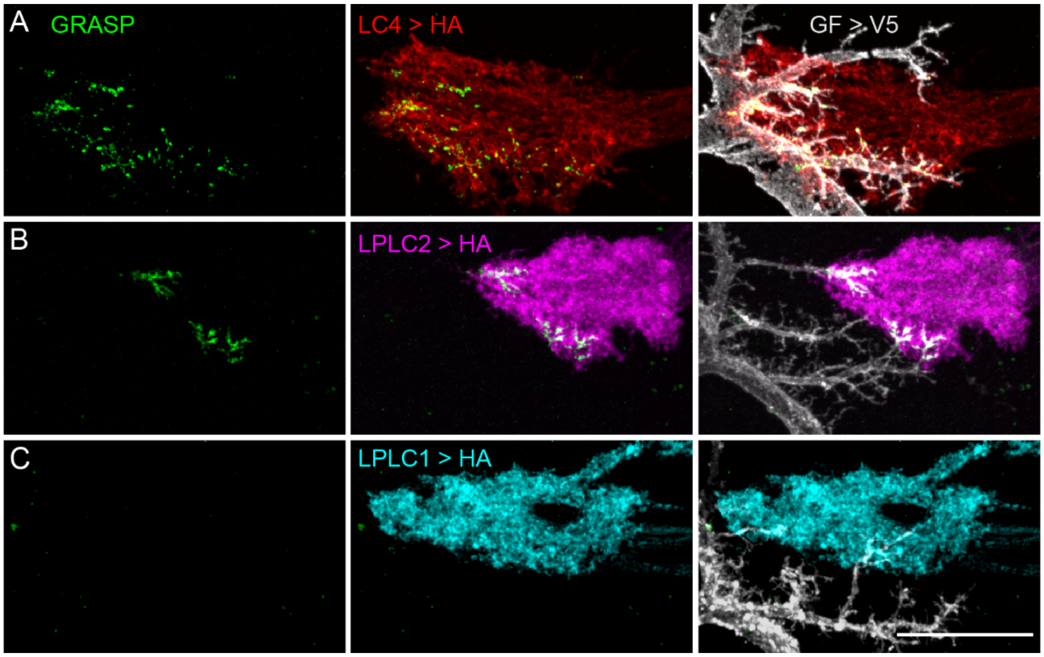

**Supplemental Figure 3. GRASP reveals putative contacts between GF and VPN**
**(A-C)** (Left) Maximum intensity projection of reconstituted GFP signal (GRASP, green) between GF and LC4 (A), LPLC2 (B) and LPLC1 (C). (Middle) Maximum intensity projection of VPN axonal membranes with GRASP signal superimposed. (Right) Maximum intensity projection of VPN axonal membranes with GRASP signal and GF (gray) superimposed. Scale bar, 20μm.

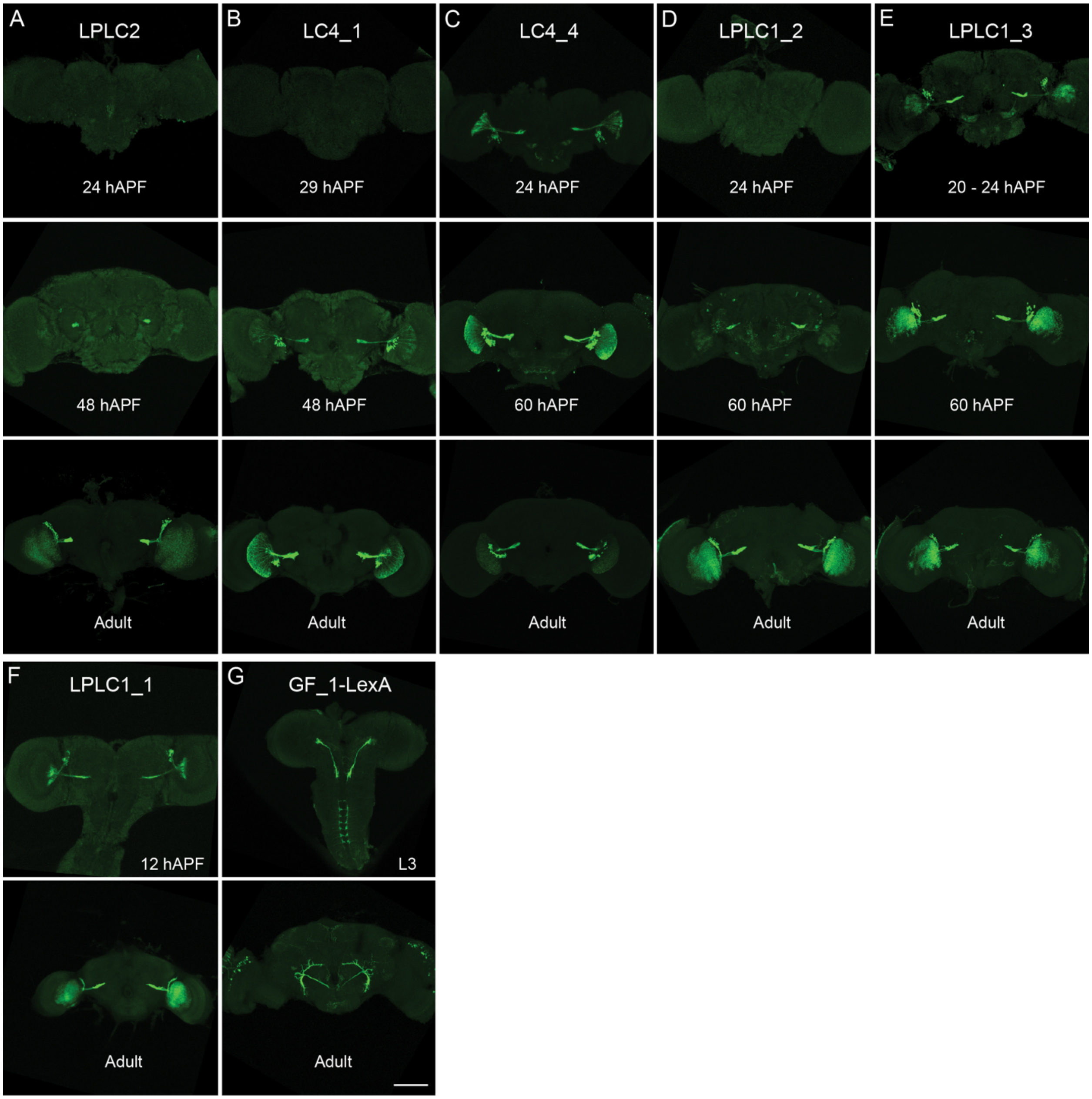

**Supplemental Figure 4. Driver line screen.**
**(A-G)** Maximum intensity projections of full brains showing developmental expression patterns (green) of VPN (Gal4 and Split-Gal4) and GF (LexA) driver lines identified and utilized within this paper. All driver lines were crossed to a smGFP-HA, smGFP-V5, or Kir2.1-mVenus reporter line. Scale bar, 50µm.

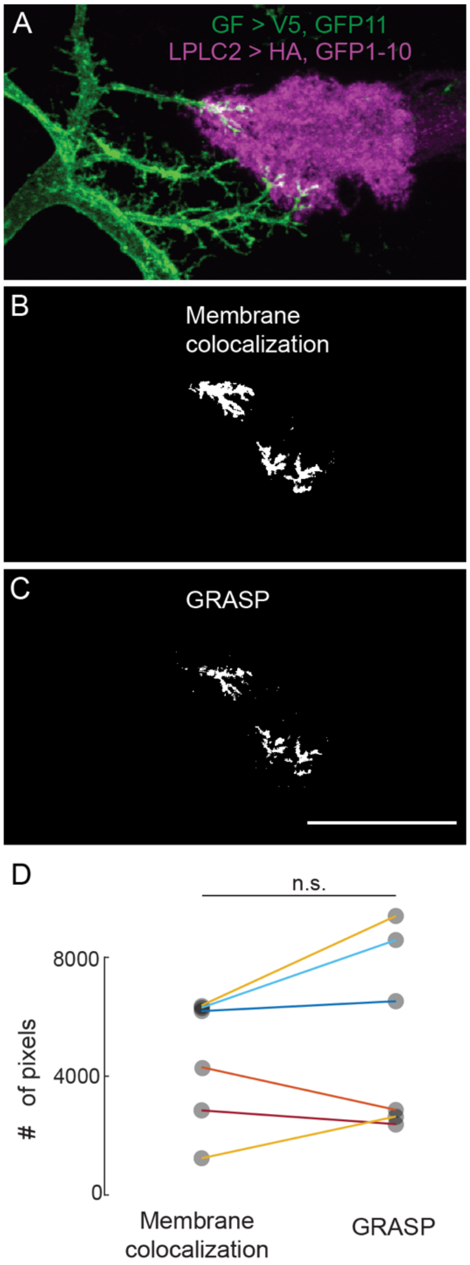

**Supplemental Figure 5. GRASP and membrane colocalization comparison**
**(A)** Maximum intensity projection image of GF (green) and LPLC2 (magenta) in adult flies.
**(B)** Z-projection of colocalized pixels (white) between GF and LPLC2 membranes.
**(C)** Z-projection of GRASP signal (white) observed between GF and LPLC2. Scale bar, 20µm.
**(D)** Quantification of colocalized pixels (B) and GRASP positive pixels (C) observed between GF and LPLC2. Paired sample *t*-test, $p = .4173$. $N = 6$ hemibrains from $\geq 3$ animals.

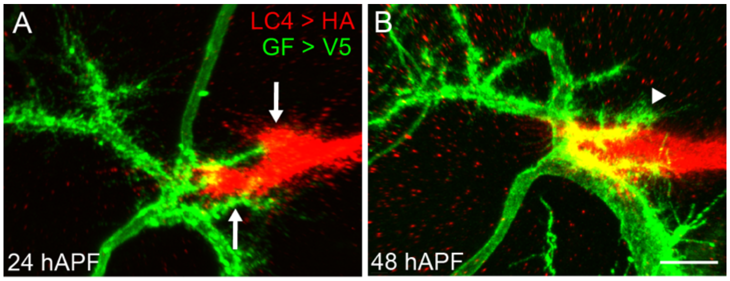

**Supplemental Figure 6. LC4 axon divergence and GF dendrite extensions during development**

**(A)** Maximum intensity projections of GF (green) and LC4 (red) at 24 hAPF. Arrows indicate dorsal (top) and ventral (bottom) division in the LC4 axon bundle.

**(B)** Maximum intensity projections at 48 hAPF. Arrowhead indicates GF dendrite extending past LC4 axons. Scale bar, 20μm.

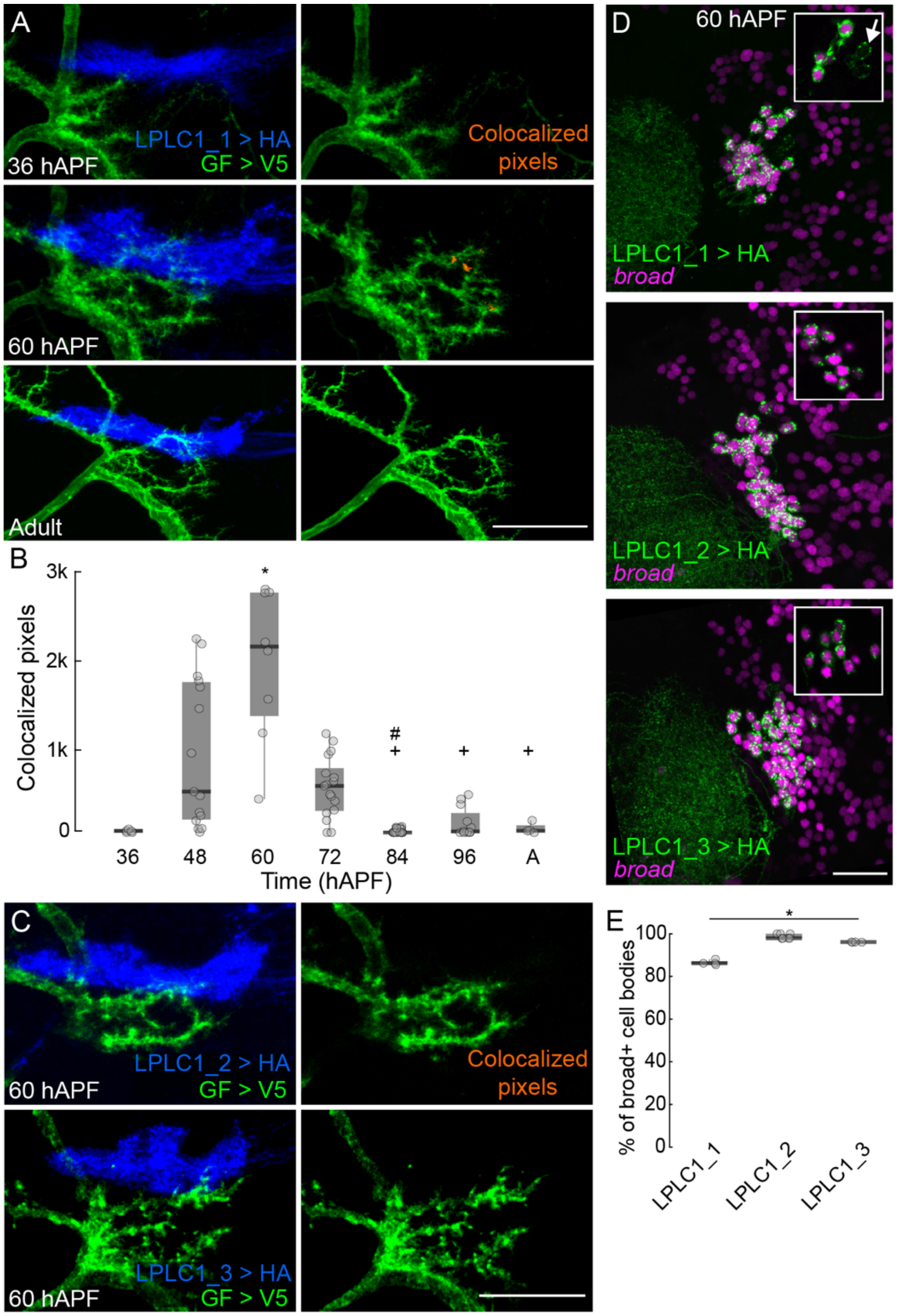

**Supplemental Figure 7. Developmental interactions between GF dendrites and LPLC1 axons**

**(A)** (Left) Maximum intensity projections of GF (green) with respect to LPLC1 (blue) at distinct developmental stages. Right, maximum intensity projections of GF with LPLC1 colocalized pixels (orange) superimposed along GF lateral dendrites. Scale bar, 20µm.

**(B)** Quantification of colocalized pixels between GF and LPLC1 in (A), an unpaired Kruskal-Wallis test was performed ($p = 1.597 \times 10^{-8}$) followed by a Dunn-Tukey-Kramer multiple comparison test post hoc. * = $p < .05$ as compared to 36 hAPF, + = $p < .05$ as compared to 60 hAPF, and # = $p < .05$ as compared to 72 hAPF. N ≥ 6 hemibrains from ≥ 3 different flies.

**(C)** (Left) Maximum intensity projection images of LPLC1-2 (top) and LPLC1-3 (bottom) driver lines with GF at 60 hAPF. (Right) Maximum intensity projection images of GF with colocalized pixels (orange) superimposed. Scale bar, 20µm. N ≥ 3 hemibrains from ≥ 3 different flies.
**(D)** Labeling specificity of LPLC1 driver lines at 60 hAPF. Maximum intensity projections of cell bodies (green) labeled in LPLC1-1 (top), LPLC1-2 (middle) or LPLC1-3 (bottom) driver lines and the LPLC1 marker broad[22] (magenta). Insets are a single z-plane, where arrows indicate cell bodies without broad expression. Scale bar, 20µm.
**(E)** Quantification of the percentage of broad-positive cell bodies in (D). N ≥ 4 hemibrains from ≥ 4 flies.

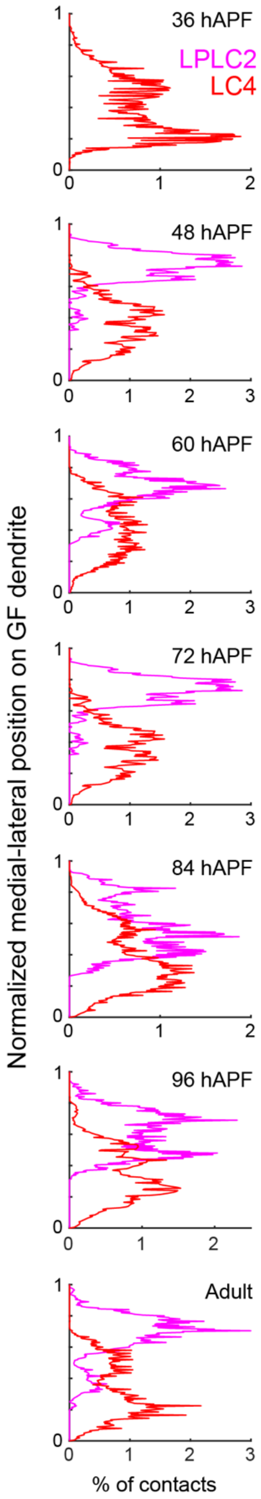

**Supplemental Figure 8. LPLC2 and LC4 contacts with GF show bias in dorsal-ventral localization**

Percentage of average VPN to GF contacts along the normalized (see methods) dorsal-ventral GF dendrite axis across development. N are as stated in Figure 2F,G.

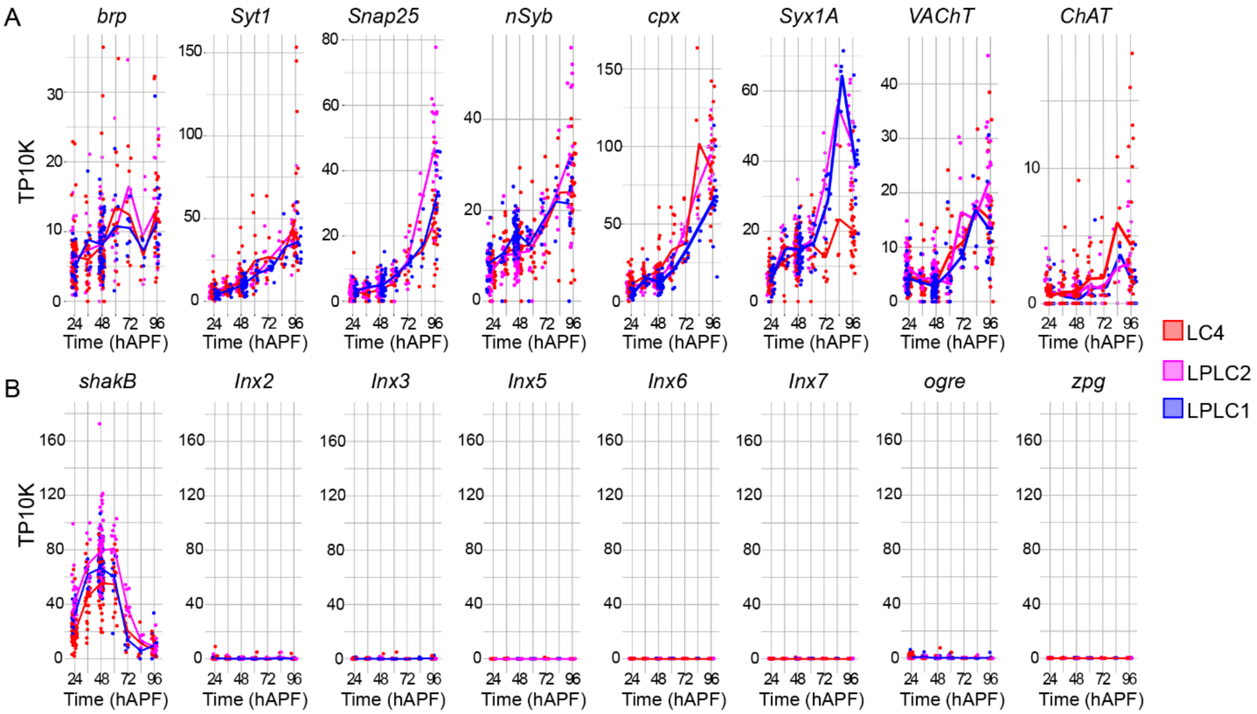

**Supplemental Figure 9. Time course of mRNA expression of presynaptic machinery and gap junction proteins in VPNs across metamorphosis**

**(A)** Expression of select presynaptic genes implicated in synaptic structure or function for LC4 (red), LPLC2 (magenta), and LPLC1 (blue) from 24 to 96 hAPF. Each point represents an individual cell, and lines represent the mean values.

**(B)** Expression of all innexins. All data are from the optic lobe transcriptional atlas[22].

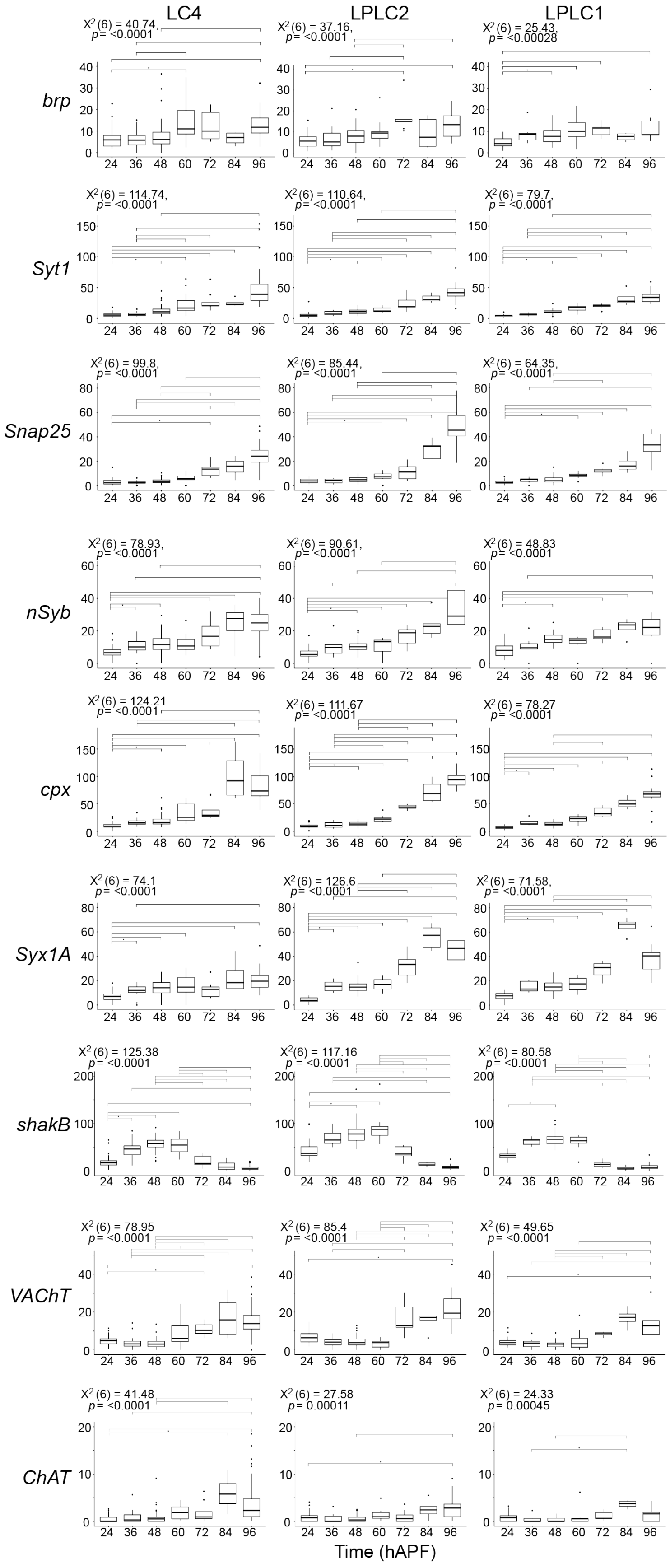
LC4
LPLC2
LPLC1
brp
X2(6) = 40.74, p= <0.0001
X2(6) = 37.16, p= <0.0001
X2(6) = 25.43, p= <0.00028
Syt1
X2(6) = 114.74, p= <0.0001
X2(6) = 110.64, p= <0.0001
X2(6) = 79.7, p= <0.0001
Snap25
X2(6) = 99.8, p= <0.0001
X2(6) = 85.44, p= <0.0001
X2(6) = 64.35, p= <0.0001
nSyb
X2(6) = 78.93, p= <0.0001
X2(6) = 90.61, p= <0.0001
X2(6) = 48.83 p= <0.0001
cpx
X2(6) = 124.21 p= <0.0001
X2(6) = 111.67 p= <0.0001
X2(6) = 78.27 p= <0.0001
Syx1A
X2(6) = 74.1 p= <0.0001
X2(6) = 126.6 p= <0.0001
X2(6) = 71.58, p= <0.0001
shakB
X2(6) = 125.38 p= <0.0001
X2(6) = 117.16 p= <0.0001
X2(6) = 80.58 p= <0.0001
VAChT
X2(6) = 78.95 p= <0.0001
X2(6) = 85.4 p= <0.0001
X2(6) = 49.65 p= <0.0001
ChAT
X2(6) = 41.48 p= <0.0001
X2(6) = 27.58 p= 0.00011
X2(6) = 24.33 p= 0.00045
24 36 48 60 72 84 96
Time (hAPF)

**Supplemental Figure 10. Statistics related to scRNA-seq analysis.**

Boxplots of within cell type and across timepoint expression comparisons for genes listed in Supplemental Figure 9. An unpaired Kruskal-Wallis test was performed and chi square values and p values are shown above each figure. Each statistically significant test was followed up with a Dunn-Bonferroni multiple comparison test, with statically significant comparisons shown in each figure. * = $p < .05$, ** = $p < .005$, *** = $p < .0005$.

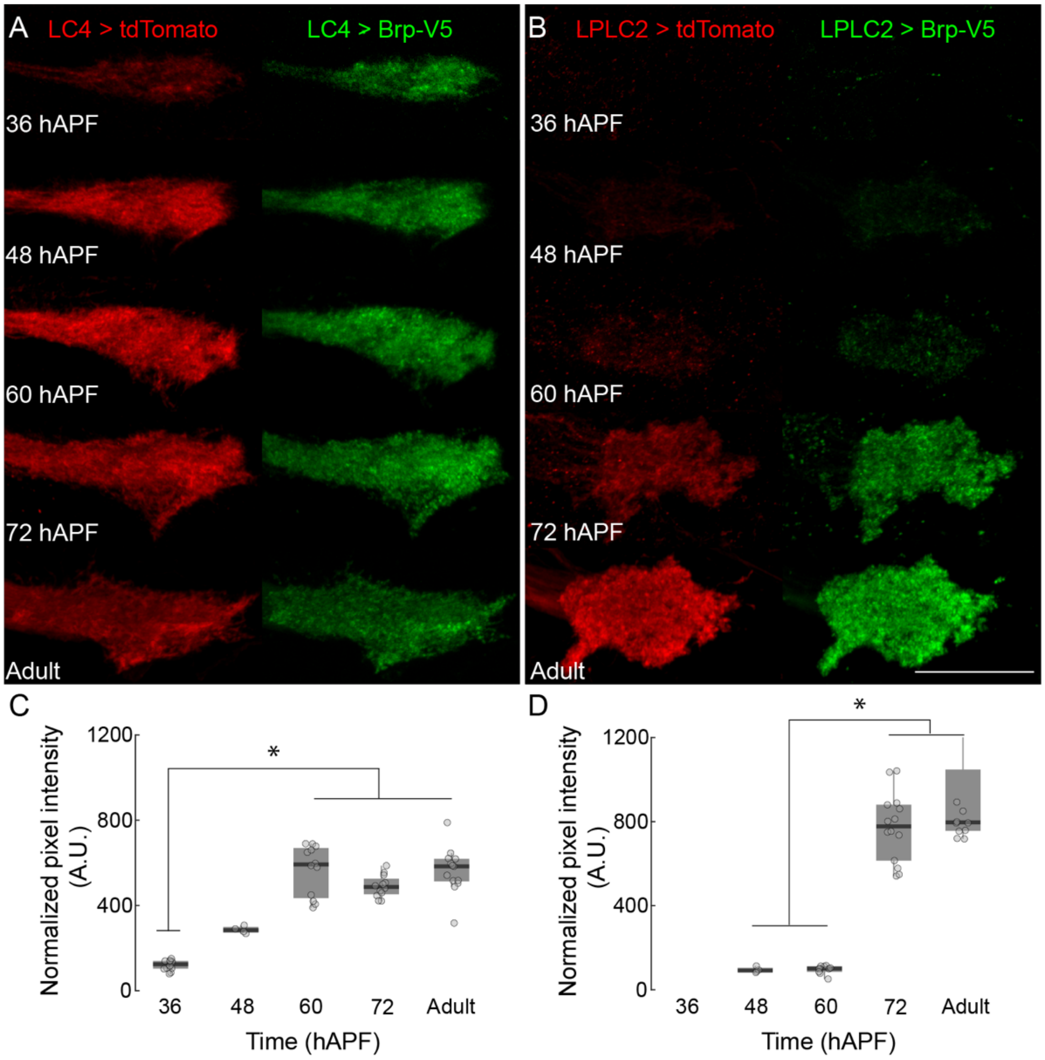

**Supplemental Figure 11. Visualization of V5 tagged bruchpilot protein in select LC populations using *smGdP-STaR***

(**A,B**) Maximum intensity projection images of LC4 (A) and LPLC2 (B) expressing tdTomato (red, left) and Brp-smGdp-V5 (green, right) under endogenous *brp* promoter at distinct times during development. Scale bar, 20µm.

**(C)** Quantification of data from A. An unpaired Kruskal-Wallis test was performed, $p = 2.423 \times 10^{-7}$, followed by Dunn-Tukey-Kramer multiple comparisons test post hoc, * = $p < .05$. $N \geq 4$ half-brains per condition from $\geq 2$ flies.

**(D)** Quantification of data from B. An unpaired Kruskal-Wallis test was performed, $p = 5.332 \times 10^{-6}$, followed by Dunn-Tukey-Kramer multiple comparisons test post hoc, * = $p < .05$. $N \geq 4$ half-brains per condition from $\geq 3$ flies.

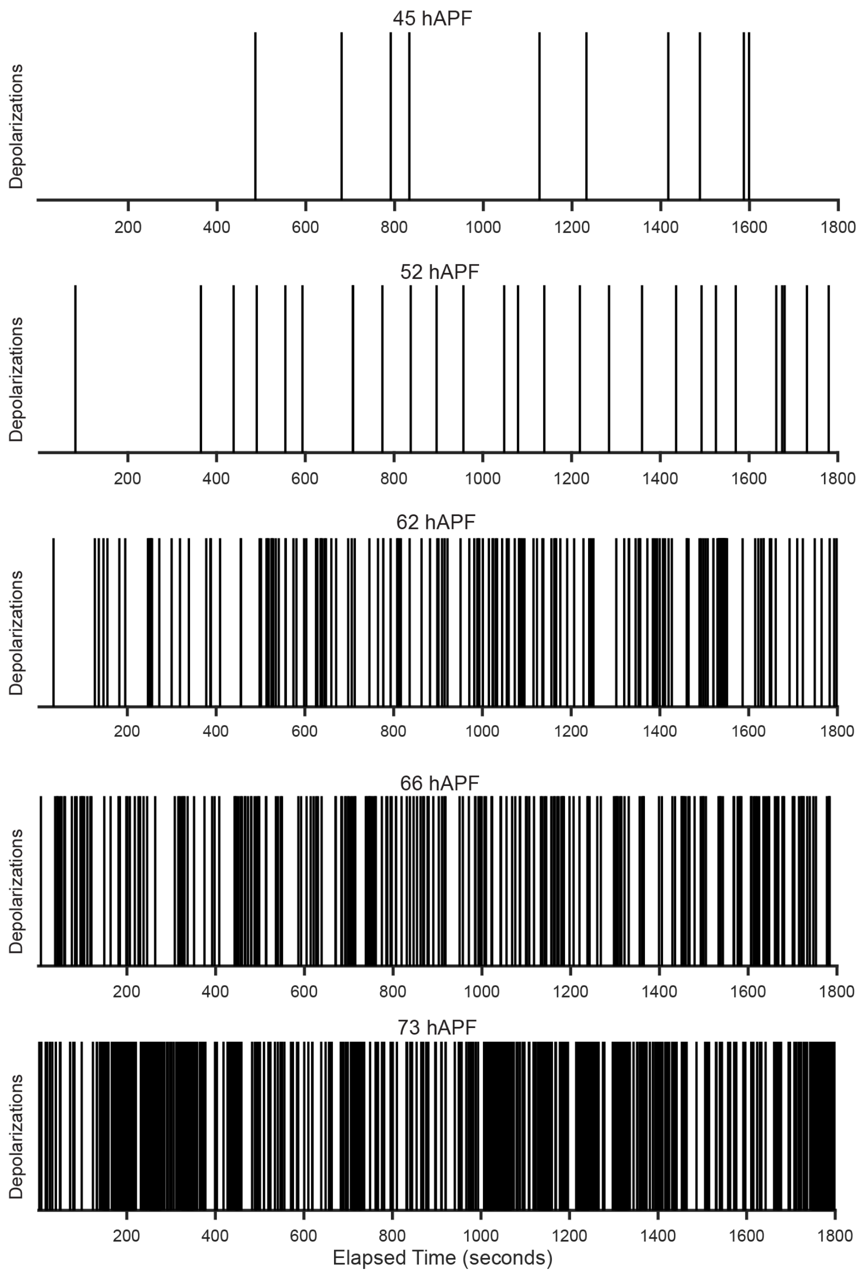

**Supplemental Figure 12. Raster plots of spontaneous activity in GF at distinct times during development**

Raster plots of each recording, each line indicates an identified depolarizing event included in our analysis.

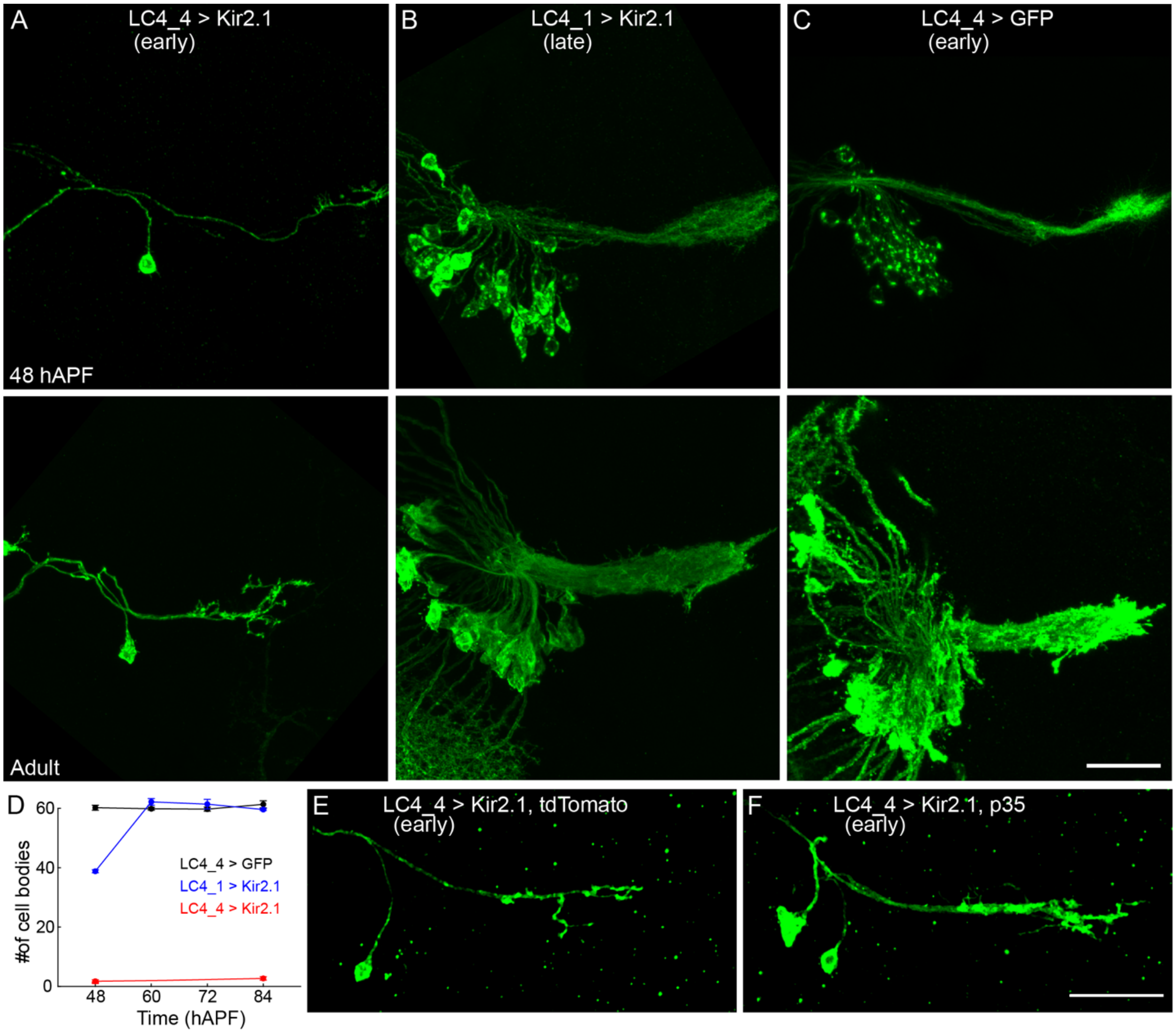

**Supplemental Figure 13. Early expression of Kir2.1 causes LC4 cell death**
(**A-C**) Maximum intensity projections of LC4 expressing Kir2.1 driven by LC4_4 (A) or LC4_1 (B), or LC4 expressing GFP driven by LC4_4 (C). Top row 48 hAPF, bottom row adult.
**(D)** Quantification of the number of LC4 cell bodies from (A-C) when expressing GFP or Kir2.1 using LC4_1 or LC4_4. N ≥ 6 hemibrains per condition from ≥ 3 flies.
(**E, F**) Maximum intensity projections of LC4 expressing Kir2.1 and tdTomato (E) or p35 (F). No significant increase in LC4 soma was observed with p35 expression (2.2 ± 0.6 tdTomato soma, 3.6 ± 0.6 p35 soma, unpaired t-test, p=0.1330, N=5-7 hemibrains from 4 flies). Scale bar, 20µm.

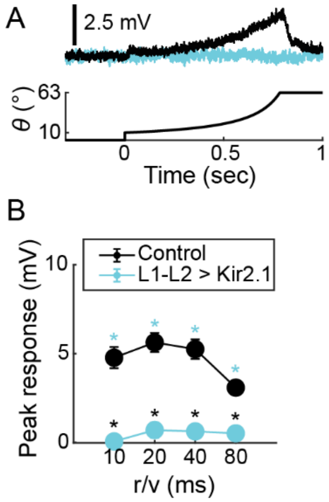

**Supplemental Figure 14. Expression of Kir2.1 in L1-L2 neurons results in a reduction of GF responses**

**(A)** Average GF responses to looming stimulus presentations of different radius to speed ratios (r/v) in control flies (black) and flies where Kir2.1 is expressed in L1-L2 neurons (cyan). Representative trace of r/v = 80ms trials.

**(B)** Quantification of peak amplitude responses to looming stimuli presentations. * = $p < .05$. N = 3-8 flies.

**Supplemental Table 1: Fly genotypes and antibodies used for each figure**

| Figure | Panel(s) | Driver Line(s) | Reporter transgene(s) | Antibodies |
|---|---|---|---|---|
| Figure 1 | A, D | *GF_1-LexA*; *LPLC2-split-GAL4*; *LC4_4-split-GAL4* | *UAS-myr::smGFP-HA, lexAop-myr::smGFP-V5* | **1° AB:**<br>mouse nc-82 (anti-Brp) (1:30)<br><br>**2° AB:**<br>647 goat anti-mouse (1:400)<br><br>Conjugate: DyLight anti-V5-550 (1:400)<br><br>Conjugate: DyLight anti-HA-488 (1:400) |
| Supplemental Figure 1 | | *GF_1-LexA*; *LPLC2-split-GAL4* | *UAS-myr::smGFP-HA, lexAop-myr::smGFP-V5* | Conjugate: DyLight anti-V5-550 (1:400).<br><br>Conjugate: DyLight anti-HA-488 (1:400) |
| Supplemental Figure 2 | B,C | *GF_1-LexA; LPLC1_1-split-GAL4* | *UAS-myr::smGFP-HA, lexAop-myr::smGFP-V5* | Conjugate: DyLight anti-V5-550 (1:400)<br><br>Conjugate: DyLight anti-HA-488 (1:400) |
| Supplemental Figure 3 | A-C | *GF_1-LexA*; *LC4_4-split-GAL4* (A); *LPLC2-split-GAL4* (B); *LPLC1_1-split-GAL4* (C) | *GRASP*<br><br>*UAS-myr::smGFP-HA, lexAop-myr::smGFP-V5* | Conjugate DyLight anti-HA-550 (1:400)<br><br>Conjugate: DyLight anti-V5-550 (1:400) |
| Figure 2 | A,D,E,H,I | *GF_1-LexA; LC4_4-split-GAL4* (D, I); | *UAS-myr::smGFP-HA,* | Conjugate: DyLight anti-V5-550 (1:400) |

| | | | | |
|---|---|---|---|---|
| | | *LPLC2-split-GAL4* (E, H) | *lexAop-myr::smGFP-V5* | DyLight anti-HA-550 (1:400) |
| Supplemental Figure 4 | A-G | *LPLC2-split-GAL4* (A); *LC4_1* (B); *LC4_4* (C); *LPLC1_2-split-GAL4*(D); *LPLC1_3-split-GAL4*(E)*;* *LPLC1_1-split-GAL4*(F); *GF_1-LexA* (G) | *UAS-myr::smGFP-HA*<br><br>*lexAop-myr::smGFP-V5*<br><br>*UAS-Kir2.1-1* | DyLight anti-HA-550 (1:400)<br><br>Conjugate: DyLight anti-V5-550 (1:400)<br><br>**$1^{o}$ AB:**<br>Chicken anti-GFP (1:1000)<br><br>**$2^{o}$ AB:**<br>488 goat anti-chicken (1:200) |
| Supplemental Figure 5 | A | *GF_1-LexA; LPLC2-split-GAL4* | *GRASP*<br><br>*UAS-myr::smGFP-HA, lexAop-myr::smGFP-V5* | Conjugate DyLight anti-HA-550 (1:400)<br><br>Conjugate: DyLight anti-V5-550 (1:400) |
| Supplemental Figure 6 | A, B | *GF_1-LexA* (A, B); *LC4_4-split-GAL4* (A, B) | *UAS-myr::smGFP-HA, lexAop-myr::smGFP-V5* | Conjugate DyLight anti-HA-550 (1:400)<br><br>Conjugate: DyLight anti-V5-550 (1:400) |
| Supplemental Figure 7 | A, C | *LPLC1_1-split-GAL4* (A, D); *LPLC1_2-split-GAL4* (C); *LPLC1_3-split-GAL4* (C); *GF_1-LexA* (A, C) | *UAS-myr::smGFP-HA, lexAop-myr::smGFP-V5* | Conjugate DyLight anti-HA-550 (1:400)<br><br>Conjugate: DyLight anti-V5-550 (1:400)<br><br>**$1^{o}$ AB:**<br>Mouse anti-broad (1:250)<br><br>**$2^{o}$ AB:** |

| | | | | |
|---|---|---|---|---|
| | | | | 647 goat anti-mouse (1:400) |
| Figure 3 | C | *LC4_4-split-GAL4*; *GF_1-LexA* | *lexAop-Brp-Short-GFP*,<br>*dlg1[4K]* | **$1^{o}$ AB:**<br>Chicken anti-GFP (1:1000)<br>**$2^{o}$ AB:**<br>488 goat anti-chicken (1:200)<br>**Conjugate:**<br>DyLight anti-V5-550 (1:400) |
| Supplemental Figure 11 | A, B | *LC4_4-split-GAL4*(A); *LPLC2-split-GAL4* (B) | *smGdP-STaR* | **$1^{o}$ AB:**<br>DsRed anti-rabbit (1:200)<br><br>**$2^{o}$ AB:**<br>568 goat anti-rabbit (1:400)<br><br>Conjugate:<br>DyLight anti-V5-650 (1:200) |
| Figure 4 | A, B, D, E, G | *LC4_4-split-GAL4*(A, B); *LPLC2-LexA*(A, B), *GF_2-LexA* (D, E, G); *LC4_1-split-GAL4* (LC4)(G) | *UAS-Kir2.1-2*(A, B, G), *UAS-Kir2.1-1*(E); *UAS-myr:GFP* (A, B, D, G); *UAS-tdTomato* (A, B, G) | **$1^{o}$ AB:**<br>Chicken anti-GFP (1:1000)<br><br>DsRed anti-rabbit (1:200)<br><br>Mouse nc82 (anti-Brp) (1:30)<br>**$2^{o}$ AB:**<br>488 goat anti-chicken (1:200)<br><br>568 goat anti-rabbit (1:400)<br><br>647 goat anti-mouse (1:400) |
| Supplemental Figure 13 | A-C, E, F | *LC4_4-split-GAL4* (LC4) (A, C, E, F); *LC4_1-split-GAL4* (LC4) (B) | *UAS-Kir2.1-1* (A, B, E, F); *UAS-tdTomato* (E); *UAS-p35* (F); | **$1^{o}$ AB:**<br>Chicken anti-GFP (1:1000) |

| | | | | |
|---|---|---|---|---|
| | | | *UAS-myr:GFP* (C) | DsRed anti-rabbit (1:200)<br><br>**2º AB:**<br>488 goat anti-chicken (1:200)<br><br>568 goat anti-rabbit (1:400) |
